# A reference metagenome sequence of the lichen *Cladonia rangiformis*

**DOI:** 10.1101/2024.12.24.630239

**Authors:** Matthias Heuberger, Carlotta Marie Wehrkamp, Alina Pfammatter, Manuel Poretti, Johannes Peter Graf, Aline Herger, Jonatan Isaksson, Edith Schlagenhauf, Rosmarie Honegger, Thomas Wicker, Alexandros G. Sotiropoulos

**Affiliations:** University of Zurich, Department of Plant and Microbial Biology, Zurich, Switzerland; University of Melbourne, Melbourne integrative genomics, Melbourne, Australia; The University of Queensland, Centre for Crop Health, Brisbane, Australia

**Keywords:** lichen, metagenome, bioinformatics, transposable element, symbiosis, *Asterochloris mediterranea*, chromosome-scale assembly, *Cladonia rangiformis*, lichens, symbiosis, microbiome

## Abstract

● Lichens are an ancient symbiosis comprising the thalli of lichen-forming fungi, their photoautotrophic partners and their microbiome. So far, they were poorly studied at the genome sequence level. Here, we present a reference metagenome for the holobiont of *Cladonia rangiformis*.

● Using long read sequences from an entire symbiotic complex, plus short read libraries from 28 additional diverse European lichen samples, we were able to separate genome sequences of 20 individual species.

● We constructed chromosome-scale assemblies of the *C. rangiformis* fungus and its trebouxioid green algal photobiont *Asterochloris mediterranea*. The genome of the fungus comprises ∼22% transposable elements and is highly compartmentalized into genic regions and large TE-derived segments which show extensive signatures of repeat-induced point mutations (RIP). We found that *A. mediterranea* centromeres are predominantly derived from two interacting retrotransposon families. We also identified strong candidates for genes that were horizontally transferred from bacteria to both alga and fungus. Furthermore, we isolated 18 near-complete bacterial genomes, of which 13 are enriched in the lichen compared to surrounding soil.

● Our study revealed that the thalli of *C. rangiformis* have a highly complex microbiome, comprising a mix of species that may include opportunists, ecologically obligate symbionts and possibly even lichen-beneficial bacteria.

## Introduction

Lichens are the symbiotic phenotype of a polyphyletic group of nutritional specialists among asco- and basidiomycetes (fungi) which acquire fixed carbon in a mutualistic symbiosis from a population of cyanobacterial or green algal cells, rarely of both; these are incorporated in the fungal thallus (Honegger 1991; Sanders 2024). Species names of lichens refer to the fungal partner, i.e. the quantitatively dominant mycobiont; the photobiont has its own name and phylogeny. Lichenisation is an ancient, ecologically successful nutritional strategy, going back at least to the lower Devonian (420 My, Honegger et al. 2013), approx. 17 % of extant fungi being lichenized (Lücking et al. 2017). Lichen thalli were defined as consortia with an unknown number of participants (Honegger 1991) or as complex ecosystems (Hawksworth and Grube 2020).

Morphologically advanced lichens have internally stratified thalli where photobiont cells are placed in optimal positions regarding illumination and gas exchange (Honegger 1986). The false reindeer lichen *Cladonia rangiformis* has a massive tubular fungal axis, the podetium, which branches abundantly in its terminal, youngest parts. The green algal photobionts (*Asterochloris mediterranea* or other *Asterochloris spp*., Moya et al. 2015) are kept in islets at the periphery of the podetia below a thin cortex. In the fruticose *C. arbuscula* large numbers of bacteria grow on the inner surface of the tubular podetium and between thick-walled, glucan-rich conglutinate hyphae of the fungal axis (Figs. 6.5.g-j in Honegger 2023). Like true reindeer lichens, *C. rangiformis* rarely reproduces sexually, pycnidia and ascomata being formed at the tip of the youngest branches. The thalli of reindeer lichens are elastic when wet, but very brittle when dry and in this state prone to fragmentation, e.g. by trampling. Fragments containing the mycobiont and photobiont and their microbiome are successfully dispersed by wind, animals or humans. Unintentional intercontinental anthropogenic dispersal of the West European *C. rangiformis* occurred (Clayden et al. 2021).

Diversity of the lichen microbiome is often assessed by using operational taxonomic units (OTUs) as proxy for species. Bacteria are typically classified based on 16S rDNA sequences, where samples with ∼98% sequence identity are defined as belonging to the same species, while ∼96% identity places them in the same genus. Eukaryotes species are routinely defined through internal transcribed spacers (ITS) sequences of rDNA, where samples of the same species usually share >98% ITS sequence identity (Janda and Abbott 2007; Edgar 2018). However, OTU counts should be considered carefully because sequence identity cutoffs may vary between studies. For example, a study on microbiomes from *Cladonia* lichens identified 79 abundant ITS OTUs, ∼81% were fungi and 18% algae (Shishido et al. 2021). However, most fungal sequences came from only four *Cladonia* species, while most green algae OTUs were from *Asterochloris mediterranea*.

The composition of microbiomes in *Cladonia* lichens seems to be generally similar. Shishido and colleagues (Shishido et al. 2021) identified 158 OTUs for bacterial 16S rDNAs, most belonging to *Alphaproteobacteria* and *Acidobacteria*. Similar findings were made in the antarctic lichen *Cladonia squamosa* (Noh et al. 2020) where most of the ∼800 OTUs belonged to the *Alphaproteobacteria* and *Acidobacteria*, with *Rhodospirillales* and *Rhizobiales* being the most abundant orders. In *C. arbuscula*, *Acetobacteraceae* closely related to the genera *Gluconacetobacter*, *Acidisphaera* and the species *Rhodovastum atsumiense* were found (Cardinale et al. 2008). However, the microbiome of *Cladoniaceae* may also vary depending on the environment (Paulsen et al. 2024), or even along the axis of single thalli (Wicaksono et al. 2020).

Studies that aim at unravelling entire lichen metagenomes are rare. The main challenge working with metagenome assemblies is that they comprise a complex mixture of thousands of sequence contigs from fungi, algae and bacteria, which have to be assigned to individual species. Genomes reconstructed from metagenomes are referred to as metagenome-assembled genomes (MAGs). Here, taxon-specific characteristics such as GC content, oligonucleotide frequencies and gene content can be used to infer to which species contigs belong. Additionally, there are specialized software such as MetaBAT2 (Kang et al. 2019) or CONCOCT (Alneberg et al. 2014) that use sequence coverage information to produce contig „bins“ which ideally represent complete MAGs.

The first lichen metagenome published was from *Umbilicaria pustulata* (Tzovaras et al. 2020), using a combination of PacBio long reads and Illumina short reads. The genome of the *U. pustulata* fungus has a size of 33 Mb, while the *Trebouxia* sp. photobiont has a 53 Mb genome. The study found that the two organisms were present at a relative abundance of fungal to algal nuclei of ∼20:1. The bacterial community was characterized through identification of sequence contigs containing 16S rDNA sequences. It was dominated by *Acidobacteriaceae* and contained bacteria similar to those typically found in lichens (e.g. *Lichenibacterium*, Pankratov et al. 2020). Very recently, 400 publicly available lichen metagenome sequences were re-examined, leading to the assembly of nearly 1000 MAGs (Tagirdzhanova et al. 2024). Genome sequences for *Cladonia* lichens are rare. We found genome assemblies for only nine *Cladonia* fungi (Table S1), four of which are assembled into (near) chromosome-scale scaffolds, while the other five are fragmented into hundreds or even thousands of sequence contigs. One reason for the fragmented assemblies may be repeat-induced point mutations (RIP), a fungal mechanism to silence transposable elements (see below). The publicly available *Cladonia* fungi genomes are 32-39 Mb in size, which is rather typical for fungi (mycocosm.jgi.doe.gov/mycocosm/home/1000-fungal-genomes). Additionally, we are aware of only four published algae genomes from the *Asterochloris* genus (Table S1.). Furthermore, a few genomes from bacteria that commonly live in *Cladonia* lichens have been assembled: *Lichenibacterium minor*, *Lichenibacterium ramalinae* and *Lichenicoccus roseus* have been grown in culture and sequenced (Pankratov et al. 2020a, b).

Considering the limited data on metagenome sequences in lichens in general and *Cladonia* in particular, we aimed at producing a reference metagenome for *C. rangiformis* that includes fungus, alga and bacteria. We report here on the composition of the chromosome-scale assemblies of the *C. rangiformis* fungus and its photobiont *A. mediterranea*. Additionally, we present MAGs for 18 bacterial species.

## Materials and Methods

### Lichen sample collection

Lichen pieces were collected from various countries around Europe including Switzerland, Greece, Iceland, France and Finland with all the sample collection details in Table S2. We also collected a soil sample next to one lichen sample and have used this particular lichen as our reference sample (i.e. GR013).

### Metagenome sequencing and assembly

Individual lichen thalli were cleaned by gently swirling them in a paper towel lined funnel under running MilliQ water. Then, they were cleaned in a petri dish filled with MilliQ water using forceps to remove all remaining dirt, animal traces and plant segments. The lichens were dried on paper towels and stored at -20°C.

High molecular weight DNA from lichen thalli was extracted using the previously published protocol for DNA extraction from wheat powdery mildew (Müller et al. 2019a) with a purification step with magnetic beads (Mayjonade et al. 2016), using used SeraSil-Mag™ silica coated superparamagnetic beads (Cytiva Europe GmbH; Freiburg im Breisgau, Germany) added for some samples (Table S4). DNA from soil samples was extracted using the NucleoSpin(R) Soil kit from MACHEREY-NAGEL (REF: 740780.10).

All samples were sequenced at the Functional Genomic Center at the University of Zurich (FGCZ) with Illumina technology using mostly 150 bp paired-end reads (Table S4). The Illumina DNA sequencing libraries were made with the Truseq PCR Free Truseq PCR Free kit to reduce the amplification bias. Additionally, the reference sample of *C. rangiformis* was sequenced with PacBio technology, creating large insert libraries, and using the PacBio Sequel system and four SMRT Cells 1M. The output reads were assembled by trying various assemblers (including SPAdes, flye, HGAP4 and megahit software) at FGCZ. The assembly with flye (metaflye mode) was one of the best and was used for further processing (Table S5). Illumina reads were then used to “polish” PacBio sequence contigs. Illumina reads for all samples were also assembled with the metaSPAdes software (v3.13.1) at default parameters. These assemblies were used for a limited set of analyses, such as species definition using ribosomal DNA (rDNA, see below). RNA was also sequenced for sample GR004 using Illumina technology (Table S4).

### Pre-processing the PacBio reference assembly

The Illumina reads of the reference genome were mapped onto the PacBio assembly using the Burrows-Wheeler Aligner (v0.7.17) (Li and Durbin 2009). We used the functions view, fixmate, sort, markdup and index from the SAMtools package to further process the mapping (Li et al. 2009; Danecek et al. 2021). The mapping was used to subsequently calculate average read coverage of individual PacBio sequence contigs.

We screened the PacBio assembly for potential chimeric contigs (i.e. sequence contigs belonging to different species that were wrongly merged). Here, we calculated GC content over a sliding window of 200 bp to identify abrupt changes in GC content along contigs in order to distinguish, for example, between segments that come from bacteria (GC content >50%) from segments that come from the fungus (GC content of <50%). Additionally, fungal genomic sequences contain regions with very low GC content as a result of RIP. These were then examined by dot plot alignments of the sequence against itself which allowed narrowing down the boundaries between regions of high and low GC content (Examples in Fig. S1). With this method we identified 9 chimeric contigs that joined fungal and bacterial segments. In two cases, the boundary could be determined precisely since it contained fungal telomeric repeats (multimers of CCCTAA) while in two cases, the boundary contains long poly C/G stretches. Chimeric contigs were then split by hand and the segments assigned either to fungal or bacterial contig groups.

### Assigning PacBio contigs to species groups

Frequencies of all 256 possible nucleotide tetramers were calculated for all PacBio sequence contigs larger than 10 kb. Shorter contigs were set aside because their tetramer frequency profiles are less representative. From this, a tetramer frequency matrix was generated which was then used for principal component analysis using R in order to separate the main species groups fungi, algae and bacteria.

Contigs shorter than 10 kb were assigned to the main species groups based on transposable element (TE) sequences. TE sequences are highly species-specific and due to their repetitive nature, can be used to assign genomic DNA segments to the respective species. Here, we first defined a reference set of large sequence contigs >300 kb that were assigned to the alga and the fungus based on tetramer frequency, the presence of known plant and fungi-specific TEs, and genes with homology to previously published *Cladonia* or *Trebouxia* genomes (see Table S1) but had no hits to contigs we defined as belonging to bacteria. This was done to exclude any contigs coming from bacteria that still may have ended up in the fungal or algal contigs. This resulted in a set of 43 high-confidence fungal and 32 algal contigs. We then used the <10KB contigs in blastn searches against the two reference datasets and identified all contigs with blastn hits >300 bp. These hits were interpreted as coming from algal or fungal TEs, respectively, and the contigs assigned accordingly to alga or fungus contig pools.

Additionally, we searched the contigs shorter than 10 kb for RIP signatures. Here, we searched for sequences that had either an overall very low GC content of less than ∼30%, and/or showed regions with abrupt changes from high to very low GC content. Of the 411 contigs identified in this approach, 403 were already assigned to the fungus in the TE search described above, which confirmed that low GC regions are mostly derived from fungal TEs. This step turned out to be largely confirmatory, since only 8 additional contigs were newly and uniquely identified through this approach to belong to the fungus.

Using tetramer frequencies for large contigs and the TE and RIP-based approaches for short contigs, we assigned 1420 contigs to the fungus and 198 to the alga pool. (see Fig. 1 and S2).

**Fig. 1.**
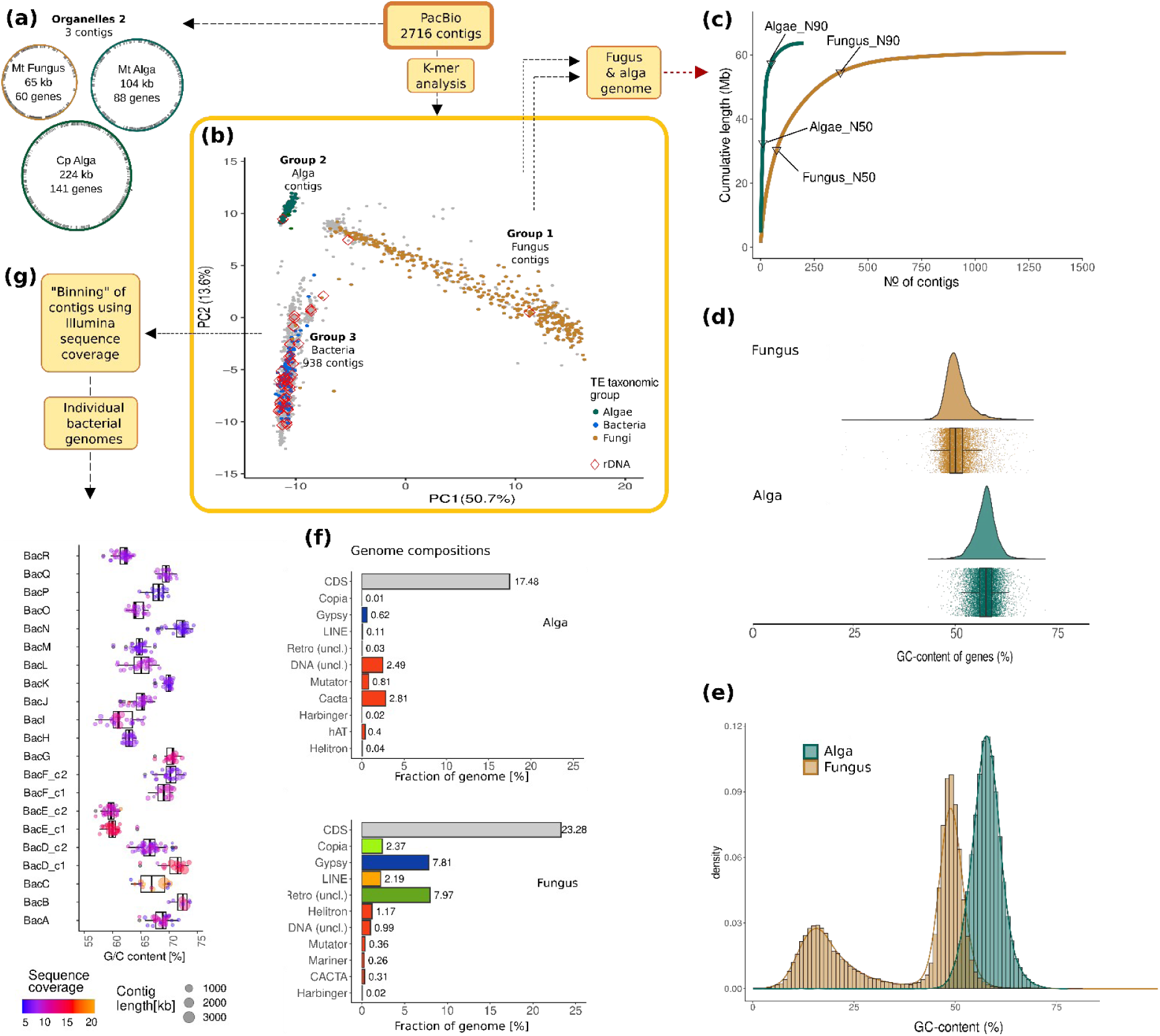
Separation of genomes from individual species from the *C. rangiformis* metagenome (a detailed workflow is shown in Fig. S2). (a) Separation of eukaryotic organelles. (b) Separation of sequence contigs into fungal, algal and bacterial contigs based on tetramer frequencies. (c) Genome coverage by sequence contigs ordered by their size. The alga fraction consists mostly of very large sequence contigs, while the fungal fraction is more fragmented. (d) GC content of coding regions (CDS) of fungal and algal genes. (e) GC content of 500 bp windows from the fungus and alga genomes. (f) Gene and TE content of the fungus and alga genomes. (g) Bacterial sequence contigs were further divided into “bins”, which ideally represent complete bacterial genomes (see Results).

### Assembly of chromosome-scale scaffolds for *C. rangiformis* and *A. mediterranea*

We used the recently published chromosome-scale assemblies of *C. squamosa* (Genbank accession GCA_947623385.2) and of an *Asterochloris* species (genbank accession GCA_963969365.1) as references to anchor PacBio contigs. Here, 1 kb segments at 10 kb intervals of the *C. squamosa* and *Asterochloris* reference chromosomes were used in blastn searches against the fungal and algal PacBio contig pool, respectively. This allowed identification of collinear segments that were then used to order the PacBio contigs into chromosomes. We were able to anchor 151 fungal and 55 algal contigs into chromosome-scale scaffolds. The 1,269 fungal and 143 algal contigs that were not included in chromosome-scale scaffolds likely represent haplotypes and/or contaminants (Fig S2, see main text).

### Gene annotation

Fungal and algal genomes were annotated using the MAKER pipeline (v. 3.01.03; Cantarel et al. 2008). Repeats were masked prior to annotation by RepeatMasker (Smit et al., 2010), employing the DFAM repeat database (Storer et al. 2021). For evidence-based annotation of the *C. rangiformis* genome, we utilized protein and transcript data from *C. grayi* (Armaleo et al. 2019), fungal hydrophobin protein sequences obtained from NCBI, and EST data from *C. rangiferina* (Junttila and Rudd 2012). Evidence-based annotation of the algal *A. mediterranea* genome utilized protein data from *Trebouxia sp. A1-2* (Tzovaras et al. 2020), and *A. thaliana* TAIR v10 proteins (Berardini et al. 2015). Additionally, a transcriptome assembly derived from *Cladonia mediterranea* RNA-seq data (comprising 219,316 contigs) was employed as EST evidence data. Initial sets of evidence-based gene models were utilized as input for the gene predictor SNAP (v2006-07-28; Korf 2004) to construct hidden Markov model (HMM) profiles. Subsequent iterative refinement through multiple rounds of SNAP training produced the ultimate gene model sets.

Completeness of genome assemblies and predicted protein-coding gene sets was assessed using BUSCO (v. 5.7.1, Simão et al. 2015; Manni et al. 2021) with the ascomycota_odb10 (2024-01-08) dataset for the fungal and the chlorophyta_odb10 (2024-01-08) dataset for the algal genome.

Reconstructed bacterial genomes (see below) were annotated with the RAST pipeline (rast.nmpdr.org) and completeness was assessed with BUSCO and CheckM (v1.1.6, Parks et al. 2015, using the lineage workflow with the reduced tree option).

### Species definition

Sequence assemblies of lichen samples were searched for contigs that contain ribosomal DNA (rDNA) genes. For fungi and algae, we extracted sequences which span short and conserved stretches from the 28S (at the 5’ end) both ITS and short stretches of 18S genes (at the 3’end). The ITS sequences were then used as queries in blastn searches against NCBI. At least the top 20 blast hits were considered because species classification of ITS sequences do not always match the morphological data (Moya et al. 2015). If blast hits were >99% identical with the sequences from the predominant species in the top blast hits, it was assumed that our sample sequence was of that species.

Bacteria were classified based on 16S rDNA. Assemblies were searched with reference 16S sequences (e.g. from *E. coli* or *Burkholderia* species). Full-length 16S genes were extracted from the assemblies and used in blastn searches against NCBI. If the top hit was >98% identical, the query was considered to be from the same species. Sequences with over ∼95% identity were considered as belonging to the same genus.

### Phylogenetic analysis

rDNA sequences were aligned with ClustalW (v2.1) and phylogenetic analyses done with MrBayes (v3.2.7a) with settings nst=6 and rates=invgamma (Ronquist et al. 2012). The Markov chain Monte Carlo (mcmc) analysis was run until the probability value decreased to under 0.01 with a sampling frequency of 10 and a burn-in of 25% of samples. Figtree (v1.4.4) (http://tree.bio.ed.ac.uk/software/figtree/) was used for visualization of phylogenetic trees.

### Reconstruction of bacterial genomes

For the further binning of the 937 bacterial contigs, we used MetaBAT2 (Kang et al. 2019), CONCOCT (Alneberg et al. 2014) and manual binning. For the manual binning, we mapped the Illumina reads of our 29 lichen samples onto our reference genome individually and calculated the sequence coverage of the contigs using the bamCoverage function from the package deepTools (Ramírez et al. 2016) from which we calculated average sequence coverage for each contig. From this, we combined values from all 29 samples into a single table using the R tidyverse library (Wickham et al. 2019). We prepared two tables using mapping data filtered by mapping quality 60 (mq60) and non-filtered reads (mapping quality 0 = mq0). From these tables we calculated correlation matrices which were clustered by hierarchical clustering utilizing the corrplot package (Version 0.92; Wei and Simko 2021) and used them for visual binning.

To be considered a genome, a group of contigs was binned together by at least two of the four approaches. Since we used the rDNA for taxonomic classification, we applied here a more stringent threshold: contigs containing rDNA had to be binned together at least three binning methods, while simultaneously being binned into the same bin which represented the majority of contigs of the genome within the respective method.

### Microbiome analysis

Illumina sequence reads were classified with the DIAMOND+MEGAN pipeline (Bağcı et al. 2021). First, they were aligned to the ncbi nr databases using diamond (v 2.1.9) blastx. Weighted binning was done using the megan (v6.25.9) command daa2rma with the weighted lca algorithm. Read information was extracted with the command rma2info. For all microbiome analyses, we only used reads that were classified at least to bacteria family level, and focused on the 10 most abundant families. Utilizing the reads2class output, we extracted the read IDs associated with these 10 families (Fig. 4A and 4b). We then subsetted these reads from the mapping files utilizing the view function from samtools (v1.13) (Li et al. 2009; Danecek et al. 2021), from which we counted reads per reconstructed bacterial genome for all 10 families. For the heat trees in Fig. 4c and 4d, comparing lichen and soil samples, we followed the “Analysis of Microbiome Community Data in R” instructions provided by the Grünwald Lab (github.com/grunwaldlab). For visualization, the node size range was set to between 1% and 5% of the graph’s width. The size range of the edges was set to 0.5% of the graph’s width. The packages metacoder (v0.3.6), taxa (v0.4.2), dplyr (v1.1.2), readr (v2.1.4) and ggplot2 (v3.4.2) were used. We obtained the necessary taxonomic classifications (kingdom, phylum, class and order) from the NCBI Taxonomy database. All bacterial families with presence in either the lichen or the soil samples were included in the figures. For clarity, labels were removed for all families with less than 1% occurrence within the sample. For calculation of these percentages, only reads from families with complete taxonomy were used. Additionally, labels for other taxonomic levels were removed, if e.g. represented families within an order all had an occurence of under 1% in the sample each.

### Identification and analysis of HT candidates

To identify HGT candidates we utilized the taxonomic classification of the Illumina short reads from the sample GR_013 we obtained from the DIAMOND+MEGAN. We extracted the names of the reads which were classified as bacteria from the reads2class output file and cross-matched them with the sam file output from mapping of the Illumina reads to our reference genome. We searched for reads that were mapped onto contigs that were integrated in fungal or algal chromosomes. Predicted genes that were covered by multiple reads were considered HGT candidates. Predicted proteins of the HGT candidates were used to search the NCBI conserved domains to determine their possible function. The HGT candidates were then used in blastp searches against the full NCBI database. If top hits were to bacterial proteins, we downloaded the top 100 sequences that were hit. To ascertain that eukaryotic homologs were more distant, we also downloaded the top 100 hits from blastp searches against the Fungi or Viridiplantae subdivision, respectively. Then, using the top 100 proteins from both searches, an alignment was produced using ClustalW (version 2.1, Larkin et al. 2007). Alignments were cropped to the aligning part of the HGT candidate and re-aligned with ClustalW. Finally, a network was created using the NeighborNet algorithm with default settings in Splitstree4 (v4.19.2) (Huson and Bryant 2006).

To date the HGT event in *A. mediterranea*, we searched for homologs in the genomes of *Trebouxia gelatinosa* (PRJNA263654) and *T. lynnae* genomes (PRJNA1123266).

## Data Availability

The scripts are deposited at https://github.com/Wicker-Lab/Cladoniaverse. Sequencing data was deposited in NCBI under the BioProject: PRJNA1168285.

## Results and Discussion

The *Cladonia* sample GR013, which was collected on the Sithonia peninsula of Chalkidiki, was chosen to generate the reference genome. The sample was whole-metagenome sequenced using PacBio long reads and Illumina short reads. The primary PacBio assembly of the *C. rangiformis* reference sample yielded 2,707 sequence contigs. Nine contigs were identified as miss-assembled hybrids and broken up accordingly (Fig. S1, methods) resulting in an assembly of 2,716 contigs with a cumulative length of 226.5 Mb. This primary assembly is a mixture of sequences from fungi, algae and bacteria which had to be assigned to individual species. To facilitate the species assignment, we also sequenced metagenomes of 28 additional lichen samples from five European countries (Greece, France, Finland, Iceland and Switzerland, Table S2) with Illumina short read technology. Since the main focus of this study were *Cladonia* lichens, 16 of the samples represent this genus, but more distantly related lichens were also included (Tables S2). The rationale for having these additional samples was to have quantitative sequence read coverage data from which the abundance of individual species in the respective samples can be inferred (i.e. species with higher abundance will be represented with higher number of sequence reads in the Illumina data). This data from the additional lichen samples proved essential in defining and isolating individual bacterial genomes in the reference *C. rangiformis* (see below).

### Separation of main groups of organisms in the *C. rangiformis* metagenome assembly

To isolate genomes of individual species we used multiple steps and criteria to separate contigs into “bins”, which ideally represented genomes of individual species. The simplified workflow is shown in Fig. 1 (more detailed in Fig. S2). First, we isolated the mitochondria and chloroplast genomes which were assembled in single contigs (Fig. 1a, Fig. S3). We then classified the remaining sequence contigs into the three main organism groups, fungi, algae and bacteria using k-mer frequencies. Here, frequencies of all 256 possible tetranucleotides were calculated for all 2,716 sequence contigs. For initial identification of the main groups, we only used contigs larger than 10 kb, because the species-specific tetramer frequencies can be poorly represented in shorter contigs. Principal component analysis (PCA) clearly segregated three distinct groups of contigs (group 1, 2 and 3, Fig. 1b, Fig. S4). Group 1 contigs showed a wide range in GC content (Fig. S4b), indicating that it represents the fungal genome. In fungal genomes we expected gene-containing regions to have higher GC content, while TE-derived sequences have very low GC content due to RIP (see below).

To assign contigs to fungi, algae and bacteria we also searched the contigs for sequences encoding canonical TE proteins such as transposase, reverse transcriptase or integrase. This confirmed group 1 contigs to represent the fungal genome, as they encode proteins with highest similarity to fungal retrotransposon superfamilies. Group 2 contigs contained numerous sequences with homology to plant TEs and thus represent the algal genome. Group 3 contigs were classified as representing bacteria genomes, because many encoded proteins with homology to typical Class 2 (DNA) transposons from bacteria, while retrotransposons were absent (Fig. 1b, S4c). Additionally, rDNA sequences identified in 41 contigs (38 bacterial, one algal and two fungal contigs) were also mapped onto the PCA (Fig. 1b. Fig. S4d). We determined the fungus to be of the species *C. rangiformis* and the alga *A. mediterranea*.

The 1,034 contigs shorter than 10 kb were assigned to the 3 main groups in a second step (Fig. S2). Here, we again took advantage of the fact that eukaryotic TEs are highly species specific. Therefore, we assigned sequence contigs that contain TE sequences to either fungus or alga based on DNA sequence homology: we used the contigs <10 kb as queries in blastn searches against the sequence contigs >10 kb that were assigned to the alga and the fungus, respectively in the previous step. Additionally, we identified fungal contigs based on signatures of repeat-induced point mutations (RIP, see methods). This allowed us to assign 847 of the contigs <10 kb to the fungus and 39 to the algal pool.

The 937 contigs classified as coming from bacteria were further subdivided separately with the goal to identify the genomes of individual bacterial species (Fig. 1c, see below).

### The genome of the *C. rangiformis* fungus shows strong signatures of repeat induced point mutations (RIP)

When analyzing sequence coverage of the 1,420 sequence contigs from fungi, we found that our sample likely consisted of two haplotypes of *C. rangiformis*, with the predominant haplotype making up at least 80% of the sample. Multiple haplotypes were previously found in *Cladonia* genome sequences (Table S1). The predominant haplotype was assembled in 151 large contigs with an Illumina read coverage of ∼220 (Fig S3, S5). We anchored these to a recently published high-quality assembly of *C. squamosa* (Genbank accession GCA_947623385.2) and produced a chromosome-scale assembly with 23 chromosomes and a total size of 34 Mb. The 1,269 contigs classified as coming from the second haplotype were collected in a separate bin and not further analyzed.

A total of 11,782 protein-coding genes were annotated on the 23 chromosomes, with 98.3% of gene models having an AED (annotation edit distance) measure below 0.5. The assembled fungal genome has a very high BUSCO (Benchmarking Universal Single-Copy Orthologs, Simão et al. 2015; Manni et al. 2021) score of >96% (Fig. S5). The *C. rangiformis* coding sequences (CDS) have a median GC content of 50.4%, similar to those of other Ascomycetes (Fig 1d, Fig. S5d). Additionally, the *C. rangiformis* genome contains ∼22% TEs. Most are retrotransposons, while only few DNA transposons were found (Fig. 1f).

Most notable, when the *C. rangiformis* genome is analyzed in 500 bp windows, it shows a bimodal GC content distribution (Fig. 1e) with gene-containing regions showing a relatively narrow peak at ∼48%, while TE-derived sequences have much lower GC content with a local peak at ∼16% (Fig. 1g), indicating high RIP activity. RIP highly efficiently silences TEs through conversion of C->T during sexual reproduction, resulting in repetitive sequences being depleted in G and C content (Cambareri et al. 1989; Mazur and Gladyshev 2018). Indeed, we found homologs of *RID* and *DIM-2*, the proposed core genes of the RIP pathway (Freitag et al. 2002). From the local minimum of the GC content distribution, we defined sequences with a GC content lower than 35% as affected by RIP (“ripped”) and those with higher GC content as “unripped”. Using this threshold, 62.5% of 500 bp genomic windows were counted as unripped and 37.5% as ripped. Similarly high levels of RIP were reported in only a few other fungi such as *Leptosphaeria maculans (syn. Plenodomus lingam)*, while most other RIP-containing ascomycetes show weaker signatures (Amselem et al. 2015). Indeed, our own analysis of 112 fungal genomes identified only 5 ascomycete species with similarly strong RIP signatures (Fig. S6).

To study to what degree genes and TEs are affected by RIP, we analyzed the GC content of genes and TEs, as well as of their flanking regions in 5’ and 3’ direction. As shown in Fig. 2, genes are practically unaffected by RIP, and their flanking sequences have in most cases roughly the same GC content. This is either due to the low-copy nature of protein coding genes, or because RIP in genes is selected against. In contrast, retrotransposons are primarily found in regions of very low GC content (Fig. 2b, 2e). This could either mean that retrotransposons specifically target “ripped” regions, or that insertions into gene-containing (i.e. “unripped”) segments are effectively selected against. Additionally, the retrotransposons themselves are mostly ripped except for a few copies which were inserted into unripped regions (Fig. 2e). Interestingly, DNA transposons are found in ripped as well as unripped regions, interspersed with genes. Furthermore, the DNA transposons themselves often show no RIP signatures (Fig. 2c, 2e, 2f, 2g and 2h). Indeed, some DNA transposon families were probably recently active, as many sequences contain newly inserted TEs which are characterized by a GC content similar to that of genic regions. Examples are the *DTM_Hades* elements shown in Fig. 2g.

**Fig. 2.**
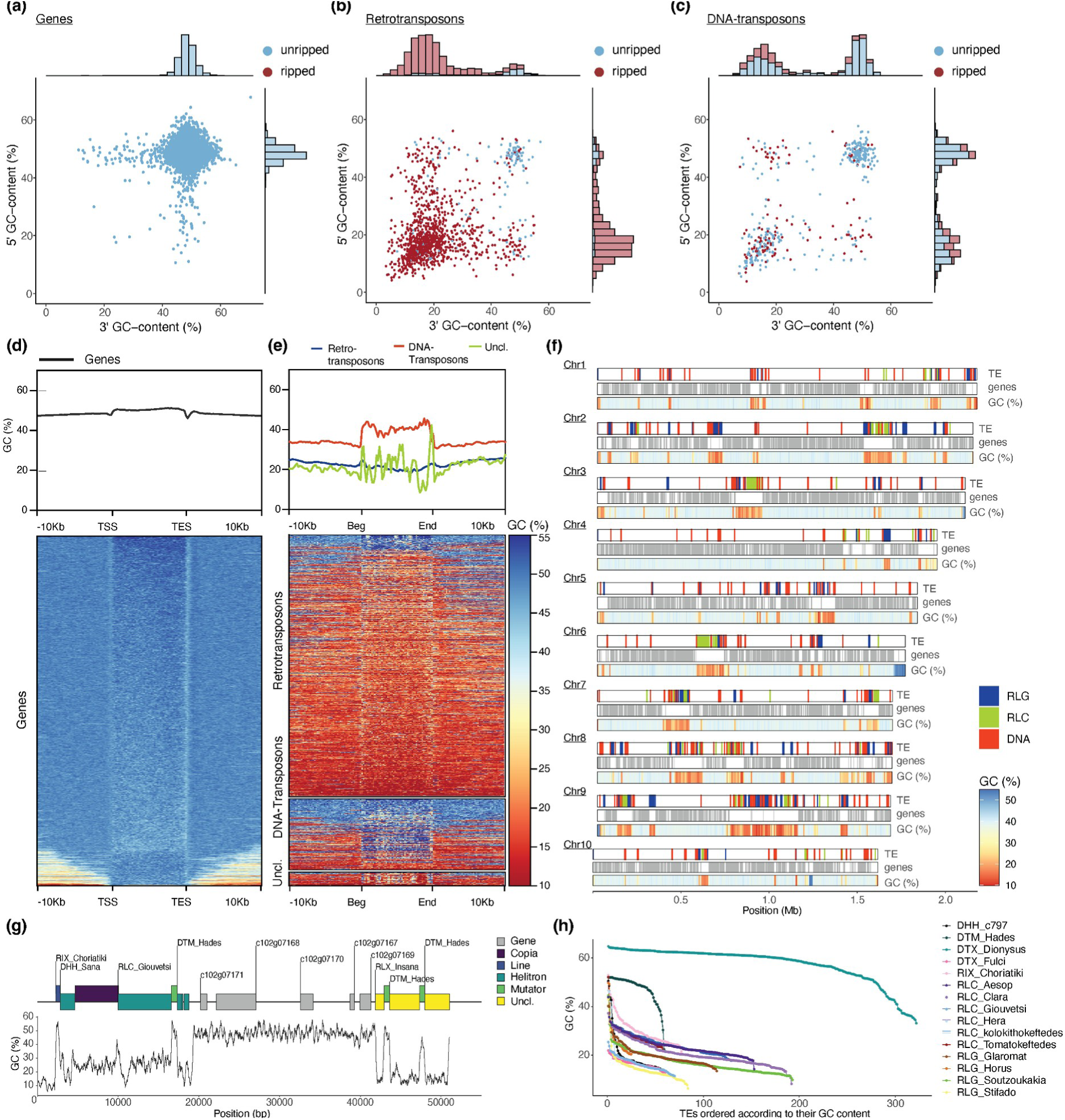
Analysis of signatures of repeat-induced point mutations (RIP) in the *C. rangiformis* genome. **(a)** GC-content of 1kb 5’ and 3’ flanking sequences of genes. Dots represent annotated genes with coloring based on whether they have a GC-content of below or above 35%. The histograms along the axes reflect the densities of dots in x and y direction. **(b)** Analogous plot for flanking sequences of Retrotransposons **(c)** Analogous plot for flanking sequences of DNA transposons. **(d)** GC-content across genes and 10kb 5’ and 3’ flanking sequence calculated in 200bp windows. Gene lengths from transcription start (TSS) to end site (TES) were normalized. The top panel shows the average values across all annotated genes, while the bottom shows a heatmap where each horizontal line corresponds to a single gene and its flanking sequences. **(e)** Analogous plot for transposable elements with retrotransposons, DNA transposons and unclassified TEs treated separately. **(f)** RIP signatures in large sequence contigs. The two top tracks show distributions of TE and genes, respectively, while the bottom track shows the GC content in 1 kb windows as a heat map. **(g) An** annotated example of a sequence contig containing ripped and unripped regions. TEs are shown as colored boxes, where nested TEs are shown raised above those into which they have inserted. Note that TEs that inserted later often have higher GC content, indicating that they went through fewer rounds of RIP. **(h)** GC content of individual copies of the 15 most abundant TE families. TE copies are ordered by their GC content from highest to lowest.

The *C. rangiformis* genome is strongly compartmentalized with regard to RIP signatures: most chromosomes have one large 100 to 500 kb RIP region, which we interpret as centromeres (example in Fig. 2f) and a few additional, somewhat smaller, RIP islands. Here again, we found DNA transposons occasionally interspersed with genes in unripped compartments, while retrotransposons are predominantly clustered in ripped regions (Fig. 2f).

The boundary between “gene space” and TEs is usually very sharp (example in Fig. 2g), indicating that TEs are precisely recognized by the RIP machinery. Additionally, there are varying levels of RIP: the majority of TE-derived sequences have a very low GC content of 10- 20%, while a fraction has an intermediate GC content of 20-40%. We interpret the latter as TEs that were more recently active and thus went through fewer RIP cycles (for example the *DHH_Sana* and *RLC_Giouvetsi* elements in Fig 2g).

Curiously, the two families of DNA transposons identified here (*DTX_Dionysus* and *DTM_Hades*) show generally much lower levels of RIP than retrotransposons, although they are found at roughly even proportions in ripped and not ripped regions (Fig. 2f). Additionally, for retrotransposon families there are only very few copies that are not ripped, presumably the most recently inserted ones, while all other copies show a rapid decline toward GC content of 10-20% when copies are ordered by their GC content (Fig. 2h). In contrast, *DTM_Hades* and *DTX_Dionysus* show the inverse GC content distribution with most copies having high GC content, while only very few underwent RIP to varying degrees (Fig. 2h). It is possible that these two TE families simply escape RIP because of their small size of ∼730 bp. Alternatively, it is possible that these two TE families were highly active very recently, and only few copies have been subjected to RIP so far.

### The centromeres of the alga *A. mediterranea* are defined by transposable elements

The *A. mediterranea* genome was assembled into much longer contigs than the fungus, despite algal contigs having a much lower sequence coverage of ∼21x. This is likely due to a low repeat content and the absence of RIP, both make contig assembly less problematic. We anchored the algal contigs to a recently published genome sequence of an *Asterochloris* species (genbank accession GCA_963969365.1) and found that we could cover the entire genome with 55 sequence contigs, with 5 contigs representing entire chromosomes. The resulting chromosome-scale assembly has a total size of 57 Mb, very similar to those coming from algae sequenced from pure cultures (Table S1). A high level of completeness is indicated by telomeric sequences at most chromosome ends (Fig. 3). The leftover 143 contigs that were classified as coming from algae are short and have lower sequence coverage (Fig. S5). They likely represent other algae species that were present at very low abundance and were not analyzed further.

**Figure 3.**
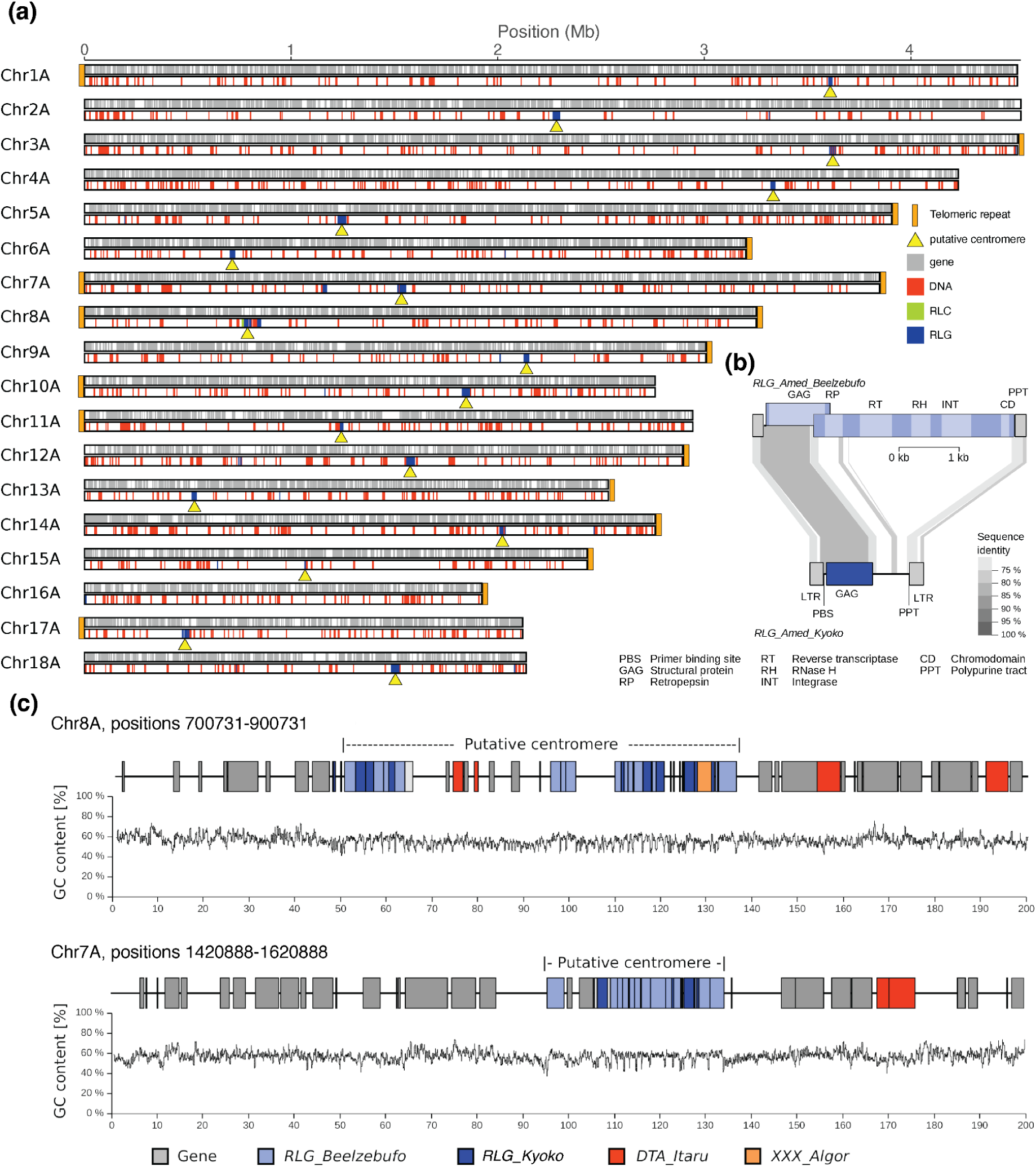
Analysis of chromosome-scale contigs in the *Asterochloris* alga genome. **(a)** Graphic representation of the 15 largest sequence contigs. The top track shows positions of annotated TEs, telomeres and putative centromeres, while the bottom track indicates positions of annotated genes. **(b)** Sequence organisation of *RLG_Beelzebufo* (top) and *RLG_Kyoko* retrotransposons (bottom). *RLG_Beelzebufo* is a putative autonomous retrotransposon, meaning it encodes all necessary enzymes for its replication, while *RLG_Kyoko* is a non-autonomous derivative which encodes only a GAG protein. Sequences which are conserved at the DNA level are connected with gray areas with darker gray indicating higher sequence identity. **(c)** Detailed annotation of two putative centromeres.

Annotation of the *A. mediterranea* genome predicted 9,107 protein-coding genes, with 88.8% of gene models having an AED measure below 0.5. BUSCO evaluation of this annotated protein-coding gene set yielded a completeness score of 90.2% ([S:88.9%,D:1.3%,F:1.3%,M:8.5%,N:1519; chlorophyta_odb10). The GC content of *Asterochloris* CDS (median 59.4%) is higher than that of the fungus (Fig. 1e) and at the higher end compared with other plants (Fig. S5b).

Only ∼8% of the *A. mediterranea* genome was annotated as TE-derived. Because no genome-wide analysis of the repetitive fraction of algae from lichens has been published so far, we focused our analysis on TE sequences. Along chromosome arms, we found predominantly DNA (Class 2) transposons interspersed with genes (Fig. 3).

Interestingly, most chromosomes contain exactly one distinct locus that is highly enriched in LTR retrotransposons, which we interpreted as centromeres (examples in Fig. 3b). Note that we use the term “centromere” here to refer to both functional centromeres and pericentromeric regions, as it is difficult to distinguish the two without epigenetic data such as CENH3 ChIP-seq. We identified two putative centromere-specific LTR retrotransposon families *RLG_Beelzebufo* and *RLG_Kyoko* (Fig. 3b). These were found in all putative centromeres (examples in Fig. 3c) but are practically absent from chromosome arms. Previous studies showed that many plant and fungi species have centromere-specific TEs (e.g. Sharma and Presting 2008; Müller et al. 2019; Ahmed et al. 2023).

The two TE families have an interesting biology: *RLG_Beelzebufo* is a putatively autonomous retrotransposon, meaning it encodes all necessary enzymes for its replication (Fig. 3b). Most importantly, it also encodes a so-called chromodomain that is fused to the integrase enzyme. Such domains are predicted to recognize the centromere-specific histone variant CENH3 and thus may target the insertion of retrotransposon copies into functional centromeres (Neumann et al. 2011; Ahmed et al. 2023; Heuberger et al. 2024). The second centromere-specific TE family, *RLG_Kyoko*, is a deletion derivative of *RLG_Beelzebufo* which lacks genes necessary for its replication such as the reverse transcriptase and integrase (Fig. 3b). We propose that *RLG_Kyoko* is cross-mobilized by *RLG_Beelzebufo* as the two fulfil the typical criteria previously described for autonomous/non-autonomous TE pairs (Wicker et al. 2022): First, the LTRs, which contain regulatory sequences, are conserved between the two, suggesting that they are likely co-expressed. Second, diagnostic motifs such as primer binding site and polypurine tract for reverse transcription are identical in the two (Fig. 3b). The situation is very similar to that in centromeres of wheat (Heuberger et al. 2024), where the autonomous *RLG_Cereba* and the non-autonomous *RLG_Quinta* retrotransposons are the main components of functional centromeres. Interestingly, this seems to have evolved independently in algae and wheat, since *RLG_Beelzebufo* and *RLG_Cereba* represent different lineages of the same superfamily containing different types of chromodomain, indicating that they acquired these domains independently.

### Isolation of bacterial genomes from the bacteria contig pool

Tetramer analysis identified a pool of 937 putative bacterial PacBio contigs, expected to represent several different bacterial species. Our aim was to group bacterial sequence contigs into „bins“, which ideally represent complete bacterial genomes. To accomplish this, we mapped Illumina sequence data from our 29 lichen samples to the 937 bacterial contigs and calculated the average read coverage for each PacBio contig across all 29 samples.

We assumed the individual bacterial species’ abundance to differ between the different lichen samples, resulting in a sample specific average sequence coverage per contig. At the same time, we expected the sequence contigs belonging to one species to have similar average sequence coverage within a sample. To group contigs into bins, we utilized the previously published softwares CONCOCT (Alneberg et al. 2014) and MetaBAT2 (Kang et al. 2019). Additionally, we produced two correlation matrices derived from two different read mapping qualities (0 and 60) for manual binning. The four approaches yielded between 36 and 47 contig bins. There was substantial overlap between contig bins generated with the four approaches, although we also frequently found differences between methods. To consolidate the results, we defined bacterial genomes as groups of contigs that are common to bins from at least two of the four approaches (Fig. S7). In total, we defined 18 bacterial genomes, comprising 7 to 56 PacBio contigs and sizes ranging from 1.04 to 5.67 Mb. A total of 377 bacterial contigs with a cumulative size of 27.7 Mb remained unassigned (Fig S2).

We assessed the completeness and quality of the retrieved genomes using BUSCO and CheckM (Fig. S8). Three genomes (BacB, BacC and BacD_chr1) had high BUSCO and CheckM completeness scores of over 85%, while seven showed medium BUSCO completeness scores ranging from 50 to 70% (BacA, BacE_chr1, BacF_chr1, BacG through BacI and BacK).

Interestingly, BacN, M, O and P show very low BUSCO and CheckM scores and are also relatively small in size (1 – 2.6 Mb). However, contigs contained in these genomes were consistently binned together in all four binning approaches (Fig. S7), which makes us confident of their assignment. We hypothesize that BacM through BacR as well as BacD_chr2, BacE_chr2 and BacF_chr2 represent secondary replicons, so called chromids. Chromids have characteristics from both bacterial chromosomes and plasmids (e.g plasmid-like replication systems, Harrison et al. 2010) and are found in about 10% of bacteria (Harrison et al. 2010). As chromids contain only few essential core genes, but rather operons for specialized metabolic pathways, this would explain the low completeness scores.

We studied the presence of the isolated bacteria genomes in our 29 lichen samples by searching for their 16S rDNA (Fig. S9). For most, we found at least rDNA from bacteria of the same genus. However, they were generally more frequent in *Cladonia* than in other samples (Fig. S9). None of them was found in all samples, which might indicate that none of them are obligate for the symbiosis. However, it is also possible that the respective rDNA was simply not covered well enough by the sequencing.

### The *C. rangiformis* microbiome comprises at least 23 bacterial species

We taxonomically classified the bacteria based on their 16S rDNA. In total, we identified 37 sequence contigs that contain 16S rDNA sequences in the 937 bacterial contigs, 32 of them were found in the 18 isolated bacterial genomes. In Bacteria, rDNA operons are found in multiple loci along bacterial chromosomes (unlike in eukaryotes where rDNA genes are usually found in a single locus). Importantly, rDNA genes coming from a single species are usually (nearly) identical due to recurring gene conversion between loci (Liao 2000). Indeed, a phylogenetic tree showed that 16S rDNA sequences coming from the same genome consistently clustered together (Fig. 4a), further validating the bacterial genome assemblies. In total, we estimate that our *C. rangiformis* metagenome assembly contains sequences from at least 23 species (Fig. 4a).

**Figure 4.**
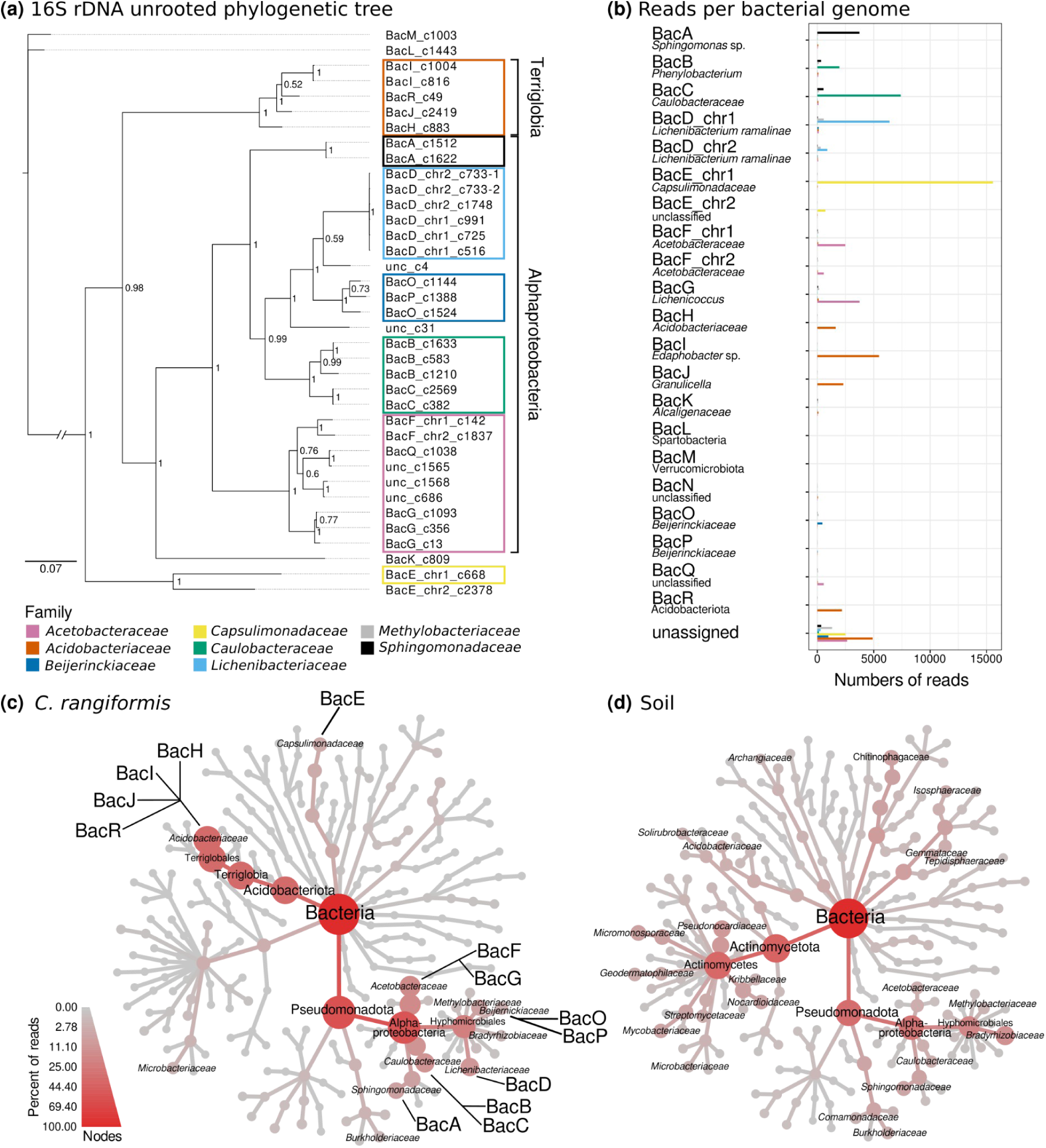
Bacterial genomes identified in the *C. rangiformis* metagenome. (**a**) Phylogenetic tree of 16S rDNA sequences. The mcmc analysis was run for 80,000 generations with a burnin value of 25%. The nodes display the probability values. Colored boxes indicate the bacterial family based on panel (b). 16S sequences retrieved from the metagenome but not assigned to a bacterial genome are labeled as unclassified (unc). (**b**) Number of reads per bacterial family and identified bacterial genome. The taxonomic classifications of the reads are indicated by the colors of the bars. Unassigned reads are the reads which were classified as one of the represented families by MEGAN but did not map to one of the retrieved bacterial genomes. Further classification is based on hits of the 16S rDNA in the NCBI database (>98% identity for species, >96% for genus and >90% for family and other taxonomic levels). (**c**) Taxonomic tree of bacterial families in the reference genome and its corresponding soil sample (**d**). The taxonomic classification was retrieved from the DIAMOND+MEGAN pipeline. Families with incidence in either the lichen or the soil samples were included in the taxonomic trees. The size of nodes and the node color represent the percentage of reads of the taxonomic levels within the respective sample. Bacterial families which were represented in more than 1% of the reads in the respective sample are labelled.

Interestingly, only four of the 18 bacteria had matches with at least 98% identity in NCBI, a commonly used cutoff for bacteria to be classified as the same species, indicating that our reference sample contains mainly poorly described bacteria. Nine of our retrieved genomes belong to the class *Alphaproteobacteria*. They could be further classified into families such as *Acetobacteraceae*, *Caulobacteraceae* and *Sphingomonadaceae* (Fig. 4a and 4b). BacN which might represent a chromid does not contain an rDNA sequence and therefore remained unclassified.

Of particular interest was BacD, which has a main chromosome (chr1) and a putative chromid (chr2). We are confident that the two chromosomes belong to the same species as all six 16S rDNA sequences found on chr1 and chr2 (3 on each) are identical (Fig 4a). Its 16S rDNA is 99% identical to that of *Lichenibacterium ramalinae* which was isolated in the subarctic region of Russia (Pankratov et al. 2020a). Interestingly, the strain present in our sample is extremely similar to the previously published one, with 41% genes being >90% identical. It is intriguing to find such closely related bacteria strains in completely different climate zones. Additionally, we found bacteria of the genus *Lichenibacterium* in 19 of our 29 lichen samples (Fig. S9). Furthermore, the published *L. ramalinae* does not contain a chr2 homolog. In our assembly, chr2 had consistently lower Illumina read coverage across *Cladonia* samples (Fig. S7), suggesting it may be present in only a fraction of the bacteria.

Bacterial genome G was classified to belong to the genus *Lichenicoccus* due to >96% identity to *Lichenicoccus roseus* (Table S6), a bacterial species isolated from *C. arbuscula* and *C. stellaris* lichens from Russia (Pankratov et al. 2020b). We found bacteria of the same genus BacG in 14 of our 16 *Cladonia* samples from north to south Europe (Fig. S9), indicating that they are a common part of lichen thalli.

Bacteria I, J and R belong to the genera *Edaphobacter* (BacI) and *Granulicella* (BacJ, BacK), which were previously described as soil bacteria adapted to arctic climate (Koch et al. 2008; Rawat et al. 2013). Our findings now indicate that they are common in *Cladonia* lichens across Europe, as we found bacteria of these genera in most of our samples (Fig. S9).

In addition to isolating bacterial genome sequences, we assessed microbiome diversity by taxonomically classifying the Illumina raw reads using the DIAMOND+MEGAN pipeline. Approximately 100,000 reads could be classified at minimum to the bacterial family level, and about 90% of them were from eight families (Fig. S10). Most of these reads mapped to our 18 bacterial genomes with only about 10% mapping to unassigned bacterial contigs (Fig. 4b). The read mapping validated the bacterial genome assemblies as all of them had predominantly reads from a single family mapped to them (Fig. 4b). Additionally, the classification using MEGAN was consistent with the phylogenetic analysis of the 16S rDNA sequences (Fig. 4a and 4b). About 4,800 reads classified as *Acetobacteriaceae* were not assigned to a genome. We therefore assume the presence of a third bacterial species from the family *Acetobacteriaceae*.

### The *C. rangiformis* microbiome differs strongly from that of surrounding soil

To gain insight which bacteria may be specific to the lichen symbiosis, we compared the microbiome of our lichens with a soil sample collected in the vicinity of the lichen thalli. The profile of bacterial families present in both samples differs considerably (Fig. 4c, 4d). In comparison to *C. rangiformis*, the soil comprises a wide variety of different bacterial families, most prominent are representatives from the class of Actinobacteria (Fig. 4d). Within the phylum Pseudomonadota, the most prevalent are Alphaproteobacteria and in particular the family *Bradyrhizobiaceae*. In *C. rangiformis* we found multiple families with comparable representation, among them *Acetobacteraceae* and *Caulobacteraceae* (Fig. 4c).

In addition, we found a strong enrichment of *Acidobacteriaceae* in the lichens in comparison to the corresponding soil. Out of the bacterial families that were found in lichens and soil, the families *Acidobacteriaceae*, *Acetobacteraceae* and *Caulobacteraceae* had the highest abundance in absolute read numbers in the *C. rangiformis* lichen samples (Fig. S11). However, the bacterial families *Capsulimonadaceae*, *Beijerinckiaceae* and *Acidobacteriaceae* are most enriched in lichens when sequence read numbers of lichen and soil are compared (Fig. S11). Most abundant with respect to absolute read counts in the soil are representatives of the families *Bradyrhizobiaceae* and *Pseudonocardiaceae* (Fig. S10), the latter also showing one of the highest relative enrichments in the soil together with the families *Mycobacteriaceae*, *Nocardioidaceae* and *Micromonosporaceae* (Fig. S12).

Especially noteworthy is the presence of representatives of *Lichenibacteriaceae*, *Lichenihabitantaceae* and *Halomonadaceae* exclusively in the *C. rangiformis* lichen, but with no incidence in the sampled soil. Furthermore, most of the taxonomic groups for which we could assemble bacterial genomes are strongly enriched in the lichen sample compared to the soil (Fig. 4c), indicating that their presence in the lichen is not coincidental. Further investigations must show whether some of these bacterial taxa are ecologically obligate symbionts of lichen thalli.

### Evidence for horizontal gene transfer between bacteria and algae and fungus

Because of the close physical proximity of the symbiotic partners in lichens, it is generally suggested that horizontal gene transfer (HGT) between them may be frequent, particularly between prokaryotic partners and fungi or algae. To identify putative HGT events, we searched for annotated genes in the *C. rangiformis* fungus and *A. mediterranea* algal chromosomes where Illumina reads mapped, that had been classified by MEGAN as coming from bacteria (Fig. S13). In this way, we identified a very strong HGT-candidate gene in *A. mediterranea* (Fig. S13b). The gene *GR013A_c05g006500* encodes a protein that shows much higher similarity to bacterial proteins from NCBI (average 93%) than to algal proteins (average ∼50%, Fig. S14). In fact, one of the closest homologs comes from *Lichenicoccus roseus* (TLU73585.1), a species for which we could reconstruct a bacterial genome (see above). The genes neighboring the HGT candidate have highest similarity to proteins from *Trebouxiophyceae* algae, indicating a single gene transfer (Fig. S13b). We emphasize that we found no indication of a sequence miss-assembly in that region, excluding the possibility of a technical artifact (Fig. S13b). Additionally, the HGT candidate has a predicted novel exon at its 5’ end that has no homology in bacterial proteins.

The HGT candidate *GR013A_c05g006500* encodes a Glutathione S-transferase (GST). GSTs are involved in a wide range of processes such as herbicide detoxification (Frear and Swanson 1970), cell signalling, plant development, and response to biotic and abiotic stresses (Roxas et al. 1997; Chen et al. 2007; Nianiou-Obeidat et al. 2017; Chronopoulou et al. 2017).

To further validate our HGT candidate we downloaded the 100 closest plant homologs from NCBI. The rationale was to determine whether plants have such homologs and whether they can be clearly distinguished from the bacterial proteins. We indeed found homologs from numerous plant taxa, including one in *A. mediterranea* (gene *GR013A_c02g008250*). We then produced a NeighborNet network that also included the top 100 bacterial homologs deposited at NCBI. The resulting network shows that the HGT candidate *GR013A_c05g006500* clusters with the bacterial proteins, while the ancestral plant homolog *GR013A_c02g008250* from *A. mediterranea* groups with other plant homologs (Fig. 5). Additionally, we did not find a *GR013A_c02g008250* homolog in published genomes of *Trebouxia* species. We conclude that plants generally have ancestral GST, but that *GR013A_c05g006500* was horizontally transferred possibly from a *Lichenicoccus* relative into *A. mediterranea* after the *Asterochloris* lineage separated from the *Trebouxia* lineage.

**Figure 5.**
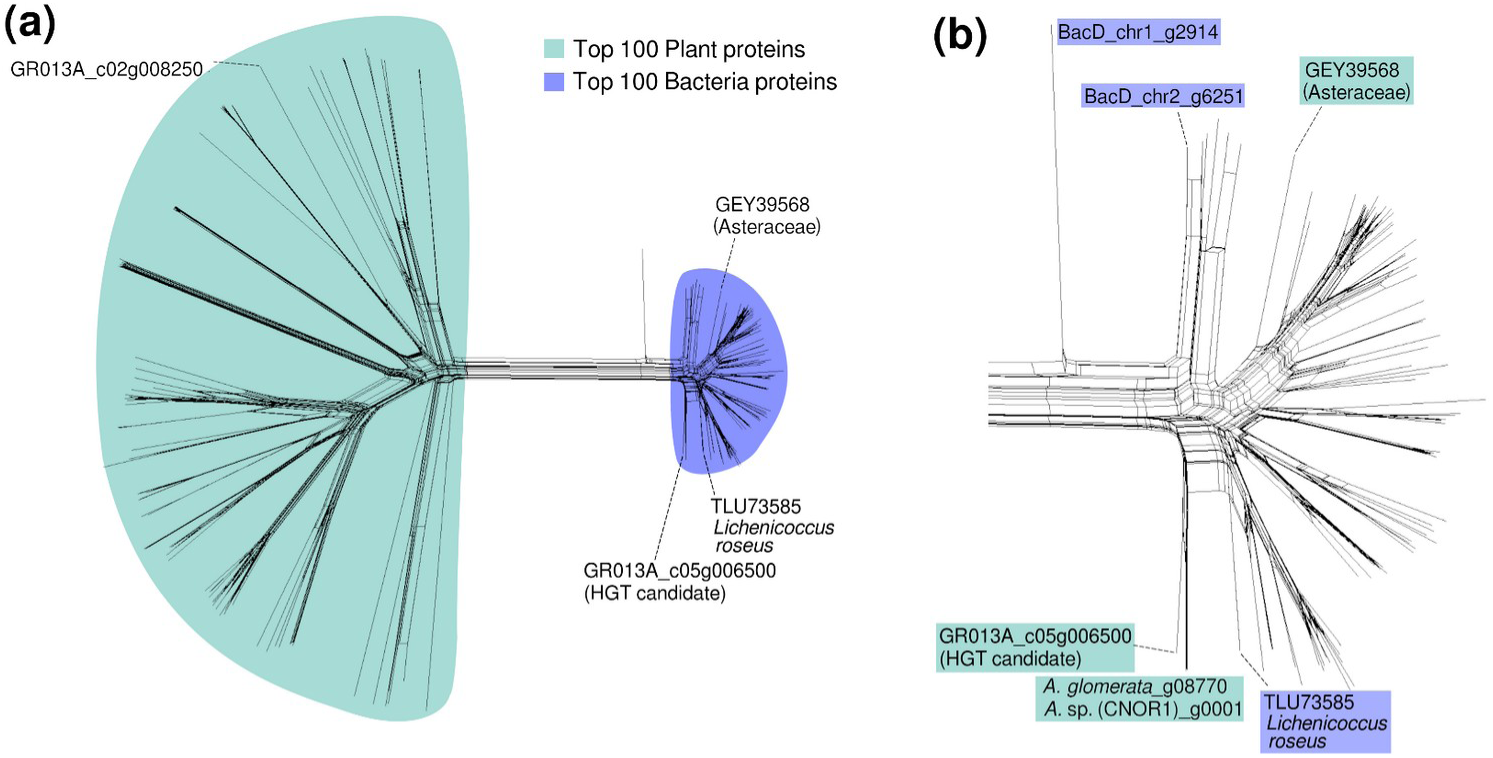
Split network of Glutathione S-transferase (GST) proteins with the HGT candidate *GR013A_c05g006500* and identified plant and bacterial homologs calculated with NeighborNet method. **(a)** Complete network with the top 100 plant and bacterial homologs. HGT homologs from Viridiplantae are marked in green, those from bacteria are shown in blue. **(b)** Zoom-in on the network of the 100 top bacterial homologs. The HGT candidate and its homologs from two publicly available *Asterochloris* genomes are shown in green. An additional homolog found in *Asteraceae* likely represents an independent HGT event. The closest bacterial homolog from NCBI (TLU7385) and those found in our bacterial genome BacD are shown in blue.

Surprisingly, the closest plant protein to GR013A_c05g006500 deposited at NCBI comes from the Dalmatian chrysanthemum (*Tanacetum cinerariifolium*), a flowering plant of the Asteraceae family. This protein also clusters with bacteria homologs, strongly indicating that this type of gene was transferred at least twice independently from bacteria to plants (Fig. 5).

Following the same method, we also identified two putative HGT events from bacteria into the fungus (Fig. S14, S15, S16). However, the network tree indicates that both HGT events must have been ancient since both HGT candidates cluster close to bacteria, but together with other fungal proteins from several different taxonomic groups (Fig. S15, S16). The two genes encode a methyltransferase domain-containing protein (*GR013F_c05g005660*), and a proline iminopeptidase-family hydrolase (GR013F_c21g001150).

## Discussion/Conclusion

To date, studies on lichen metagenomes are still rare, likely due to the complex task of isolating and analyzing genomes from multiple and diverse species. A recent study published the metagenome of the lichen *U. pustulata* (Tzovaras et al. 2020). However, that study focused on the assembly of fungal and algal genomes, while analysis of bacterial genomes was limited to rDNA. In contrast, 400 publicly available lichen metagenome sequences were re-examined, leading to the assembly of nearly 1000 MAGs (Tagirdzhanova et al. 2024), most of them from bacterial species.

Because of rapid technological development, the number of lichen metagenomes will grow in the future. We, therefore, considered it timely to assess and combine multiple different strategies to assemble high-quality MAGs for the fungus, alga and bacteria. Here, we want to emphasize a few particularly successful approaches: At a basic level, k-mer analysis proved highly effective in separating the main organism groups: fungus, alga and bacteria. This method also worked for much more fragmented short read assemblies of lichen metagenomes (Puvar et al. 2020). Additionally, because of their species-specificity and repetitive nature, TE sequences were highly effective to distinguish between plant and fungal sequences, and in some cases even between bacteria. To assemble chromosome-scale scaffolds for *C. rangiformis* and *A. mediterranea* we also took advantage of recently published genome assemblies. Although from different species, chromosomes were sufficiently similar to anchor our sequence contigs. Furthermore, for assemblies of bacterial genomes, considering sequence read coverage was essential. Here, it proved particularly useful that we had metagenome data from additional various lichens. Our combined use of the correlation matrices as well as the software CONCOCT (Alneberg et al. 2014) and MetaBat (Kang et al. 2019) allowed reliable reconstruction of bacterial genomes, while use of only one of the approaches may not be sufficient. Ultimately, we succeeded in assigning ∼80% of the ∼220 Mb assembly to individual species.

Due to limited resources, the initial assembly was done on PacBio Sequel technology. It is conceivable that use of the latest long read technology combined with chromatin conformation capture (Hi-C), may produce less fragmented meta-genome assemblies. Nevertheless, our chromosome-scale assemblies are of comparable quality as recently published algal and fungal genomes that were produced from pure cultures (e.g. *C. grayi* and *A. glomerata*, Armaleo et al. 2019). However, while organelles assembled into single contigs, our bacterial MAGs are still fragmented to some degree.

The chromosome-scale assemblies allowed detailed insight into the organization of gene-containing and repetitive fractions of *C. rangiformis* and *A. mediterranea*, analyses that, to our knowledge, have not been done in lichens so far. We found both genomes to be highly compartmentalized regarding TE sequences. In *A. mediterranea*, we identified a pair of centromere-specific TEs that had strikingly resembling counterparts in grasses (Heuberger et al. 2024) where autonomous *RLG_Cereba* and non-autonomous *RLG_Quinta* retrotransposons are the main components of functional centromeres. Interestingly, the two systems must have evolved independently in algae and grasses, since *RLG_Beelzebufo* and *RLG_Cereba* represent different evolutionary lineages and contain different types of chromodomains, indicating that they acquired these domains independently.

The *C. rangiformis* fungus showed very strong RIP signatures, levels of which have been found in only a few fungi so far. RIP is proposed to occur during sexual reproduction (Cambareri et al. 1989). The fact that we find only very few “unripped” TE sequences could mean that *C. rangiformis* frequently undergoes sexual recombination, despite clonal propagation being an important mechanism for the spread of symbiotic “propagules” (Yahr et al. 2006). Alternatively, *Cladonia* fungi may have TE silencing pathways additional to RIP. Interestingly, we also identified putative HGT events from bacteria into the eukaryotic partners. However, it is notoriously difficult to prove HGT, especially if it occurred a long time ago and/or the prokaryotic gene donor is not known. Indeed, while some studies reported HGT (Armaleo et al. 2019), others found no evidence (Tzovaras et al. 2020). We were able to detect the origin of the GST which we regard as a likely horizontally transferred gene by, amongst others, relying on the annotation of proteins in the NCBI database. If proteins originating from assemblies of lichen (meta-)genomes are annotated as such and deposited in the NCBI database, a horizontally transferred protein will potentially have a higher similarity to proteins from other lichens than to the donor of the HT sequence, for example bacteria, when using the blastp search against the NCBI database. We, therefore, expect that the detection of HGT events becomes increasingly difficult as more metagenomic resources for lichens become available.

Our study was particularly informative with regard to the microbiome. The 18 bacteria species for which we assembled genomes were found at much lower abundance (or not at all) in surrounding soil, indicating they specifically proliferate in the lichen thallus. There are likely additional, less abundant species which we did not capture with our sequencing depth. Nevertheless, a recent study reported bacteria of the families *Acidobacteriaceae*, *Acetobacteraceae*, *Beijerinckiaceae* and *Sphingomonadaceae* to occur most frequently in lichens (Tagirdzhanova et al. 2024). These four families were also present in our 18 bacterial genomes. Additionally, all 18 were also found at least to some level in our 16 *Cladonia* samples coming from all over Europe. This indicates that the bacterial genomes isolated here are common and abundant cohabitants of lichens. Furthermore, we found two bacterial genera (*Lichenicoccus* and *Sphingomonas*) that were described in a previous study (Tagirdzhanova et al. 2024).

Particularly surprising were the presence of *Edaphobacter* and *Granulicella* species, previously described as soil bacteria adapted to arctic climate (Koch et al. 2008; Rawat et al. 2013). We found them in consistent abundance in samples from north to south Europe. The two genera seem to be adapted to low carbon concentrations and metabolizing complex polysaccharides (Koch et al. 2008; Rawat et al. 2013). These bacterial inhabitants of lichen thalli might mobilize carbohydrates from the glucan-rich fungal cell walls for their own nutrition. At this point it is still not clear which roles the bacteria play in the symbiosis and whether there are additional species which we did not detect due to low abundance. Contrary to our expectations, we found none of the marker genes associated with nitrogen fixation, as reported in a recent study (Tagirdzhanova et al. 2024), indicating that nitrogen fixation is not a typical feature of the microbiome of *C. rangiformis*. Moreover, we also found none of the previously described lichenicolous fungi (Pino-Bodas and Stenroos 2021). In plants mycorrhiza helper bacteria (Frey-Klett et al. 2005) and a wide range of other plant-beneficial bacteria have been characterized (Vandenkoornhuyse et al. 2015; Trivedi et al. 2022). With high probability some representatives of the microbiome of lichen thalli are lichen-beneficial bacteria; their potential roles in the symbiosis remain to be explored.

Our *C. rangiformis* reference metagenome provides a glance at species diversity, but much broader research is needed for our understanding of the full complexity and interplay of the different taxa in the thalli of *Cladonia* species.

## Acknowledgements

This work was supported by the Swiss National Science Foundation grant 310030_212428, University of Zurich core funding and the University Research Priority Program of Evolution in Action of the University of Zurich.

## Author contributions

TW and AGS conceived the project. AGS, CMW, AP, MP, AH, JI and TW collected samples. AGS, CMW, AP and MH extracted DNA. MH, CMW, AGS, TW, ES, JPG, MP and AP conducted bioinformatic analyses. MH, CMW, AGS, ES, TW and RH wrote the initial manuscript with revisions from MP, AH, JI. MH and CMW contributed equally to this work.

## References

1. Ahmed HI, Heuberger M, Schoen A, et al (2023) Einkorn genomics sheds light on history of the oldest domesticated wheat. Nature 620:830–838. 10.1038/s41586-023-06389-7

2. Alneberg J, Bjarnason BS, De Bruijn I, et al (2014) Binning metagenomic contigs by coverage and composition. Nat Methods 2014 1111 11:1144–1146. 10.1038/nmeth.3103

3. Amselem J, Lebrun MH, Quesneville H (2015) Whole genome comparative analysis of transposable elements provides new insight into mechanisms of their inactivation in fungal genomes. BMC Genomics 16:1–14. 10.1186/S12864-015-1347-1

4. Armaleo D, Müller O, Lutzoni F, et al (2019) The lichen symbiosis re-viewed through the genomes of Cladonia grayi and its algal partner Asterochloris glomerata. BMC Genomics 20:1–33. 10.1186/s12864-019-5629-x

5. Bağcı C, Patz S, Huson DH (2021) DIAMOND+MEGAN: Fast and Easy Taxonomic and Functional Analysis of Short and Long Microbiome Sequences. Curr Protoc 1:. 10.1002/CPZ1.59

6. Berardini TZ, Reiser L, Li D, et al (2015) The Arabidopsis information resource: Making and mining the “gold standard” annotated reference plant genome. Genesis 53:474–485. 10.1002/DVG.22877

7. Cambareri EB, Jensen BC, Schabtach E, Selker EU (1989) Repeat-induced G-C to A-T Mutations in Neurospora. Science (80-) 244:1571–1575. 10.1126/SCIENCE.2544994

8. Cantarel BL, Korf I, Robb SMC, et al (2008) MAKER: an easy-to-use annotation pipeline designed for emerging model organism genomes. Genome Res 18:188–196. 10.1101/GR.6743907

9. Cardinale M, Vieira De Castro J, Müller H, et al (2008) In situ analysis of the bacterial community associated with the reindeer lichen Cladonia arbuscula reveals predominance of Alphaproteobacteria. FEMS Microbiol Ecol 66:63–71. 10.1111/J.1574-6941.2008.00546.X

10. Chen IC, Huang IC, Liu MJ, et al (2007) Glutathione S-Transferase Interacting with Far-Red Insensitive 219 Is Involved in Phytochrome A-Mediated Signaling in Arabidopsis. Plant Physiol 143:1189–1202. 10.1104/PP.106.094185

11. Chronopoulou E, Ataya FS, Pouliou F, et al (2017) Structure, Evolution and Functional Roles of Plant Glutathione Transferases. Glutathione Plant Growth, Dev Stress Toler 195–213. 10.1007/978-3-319-66682-2_9

12. Clayden S, Ahti T, Pino-Bodas R, et al (2021) First documented occurrences of Cladonia krogiana and C. rangiformis in North America. Opusc Philolichenum 20:25–36. 10.5962/p.388271

13. Danecek P, Bonfield JK, Liddle J, et al (2021) Twelve years of SAMtools and BCFtools. Gigascience 10:giab008. 10.1093/gigascience/giab008

14. Edgar R (2018) Taxonomy annotation and guide tree errors in 16S rRNA databases. PeerJ 2018:e5030. 10.7717/PEERJ.5030

15. Frear DS, Swanson HR (1970) Biosynthesis of S-(4-ethylamino-6-isopropylamino- 2-s- triazino) glutathione: Partial purification and properties of a glutathione S-transferase from corn. Phytochemistry 9:2123–2132. 10.1016/S0031-9422(00)85377-7

16. Freitag M, Williams RL, Kothe GO, Selker EU (2002) A cytosine methyltransferase homologue is essential for repeat-induced point mutation in Neurospora crassa. Proc Natl Acad Sci U S A 99:8802–8807. 10.1073/PNAS.132212899

17. Frey-Klett P, Chavatte M, Clausse ML, et al (2005) Ectomycorrhizal symbiosis affects functional diversity of rhizosphere fluorescent pseudomonads. New Phytol 165:317–328. 10.1111/J.1469-8137.2004.01212.X

18. Harrison PW, Lower RPJ, Kim NKD, Young JPW (2010) Introducing the bacterial “chromid”: not a chromosome, not a plasmid. Trends Microbiol 18:141–148. 10.1016/J.TIM.2009.12.010

19. Hawksworth DL, Grube M (2020) Lichens redefined as complex ecosystems. New Phytol 227:1281–1283. 10.1111/NPH.16630

20. Heuberger M, Koo D-H, Ahmed HI, et al (2024) Evolution of Einkorn wheat centromeres is driven by the mutualistic interplay of two LTR retrotransposons. Mob DNA 15:16

21. Honegger R (1991) Functional aspects of the lichen symbiosis. Annu Rev Plant Physiol Plant Mol Biol 42:553–578. 10.1146/ANNUREV.PP.42.060191.003005

22. Honegger R (2023) Lichens and Their Allies Past and Present. 133–183. 10.1007/978-3-031-16503-0_6

23. HONEGGER R (1986) Ultrastructural Studies in Lichens: I. Haustorial Types and Their Frequencies in a Range of Lichens With Trebouxioid Photobionts. New Phytol 103:785–795. 10.1111/j.1469-8137.1986.tb00853.x

24. Honegger R, Axe L, Edwards D (2013) Bacterial epibionts and endolichenic actinobacteria and fungi in the Lower Devonian lichen Chlorolichenomycites salopensis. Fungal Biol 117:512–518. 10.1016/J.FUNBIO.2013.05.003

25. Huson DH, Bryant D (2006) Application of phylogenetic networks in evolutionary studies. Mol Biol Evol 23:254–267. 10.1093/MOLBEV/MSJ030

26. Janda JM, Abbott SL (2007) 16S rRNA gene sequencing for bacterial identification in the diagnostic laboratory: pluses, perils, and pitfalls. J Clin Microbiol 45:2761–2764. 10.1128/JCM.01228-07

27. Junttila S, Rudd S (2012) Characterization of a transcriptome from a non-model organism, Cladonia rangiferina, the grey reindeer lichen, using high-throughput next generation sequencing and EST sequence data. BMC Genomics 13:. 10.1186/1471-2164-13-575

28. Kang DD, Li F, Kirton E, et al (2019) MetaBAT 2: An adaptive binning algorithm for robust and efficient genome reconstruction from metagenome assemblies. PeerJ 2019:e7359. 10.7717/PEERJ.7359

29. Koch IH, Gich F, Dunfield PF, Overmann J (2008) Edaphobacter modestus gen. nov., sp. nov., and Edaphobacter aggregans sp. nov., acidobacteria isolated from alpine and forest soils. Int J Syst Evol Microbiol 58:1114–1122. 10.1099/IJS.0.65303-0

30. Korf I (2004) Gene finding in novel genomes. BMC Bioinformatics 5:. 10.1186/1471-2105-5-59

31. Larkin MA, Blackshields G, Brown NP, et al (2007) Clustal W and Clustal X version 2.0. Bioinformatics 23:2947–2948. 10.1093/BIOINFORMATICS/BTM404

32. Li H, Durbin R (2009) Fast and accurate short read alignment with Burrows-Wheeler transform. Bioinformatics 25:1754–1760. 10.1093/bioinformatics/btp324

33. Li H, Handsaker B, Wysoker A, et al (2009) The Sequence Alignment/Map format and SAMtools. Bioinformatics 25:2078–2079. 10.1093/BIOINFORMATICS/BTP352

34. Liao D (2000) Gene conversion drives within genic sequences: concerted evolution of ribosomal RNA genes in bacteria and archaea. J Mol Evol 51:305–317. 10.1007/S002390010093

35. Lücking R, Hodkinson BP, Leavitt SD (2017) The 2016 classification of lichenized fungi in the Ascomycota and Basidiomycota – Approaching one thousand genera. 101639/0007-2745-1194361119:361–416. 10.1639/0007-2745-119.4.361

36. Manni M, Berkeley MR, Seppey M, Zdobnov EM (2021) BUSCO: Assessing Genomic Data Quality and Beyond. Curr Protoc 1:. 10.1002/CPZ1.323

37. Mayjonade B, Gouzy J, Donnadieu C, et al (2016) Extraction of high-molecular-weight genomic DNA for long-read sequencing of single molecules. Biotechniques 61:203–205. 10.2144/000114460

38. Mazur AK, Gladyshev E (2018) Partition of Repeat-Induced Point Mutations Reveals Structural Aspects of Homologous DNA-DNA Pairing. Biophys J 115:605–615. 10.1016/j.bpj.2018.06.030

39. Moya P, Škaloud P, Chiva S, et al (2015) Molecular phylogeny and ultrastructure of the lichen microalga asterochloris mediterranea sp. nov. from mediterranean and Canary Islands ecosystems. Int J Syst Evol Microbiol 65:1838–1854. 10.1099/IJS.0.000185

40. Müller MC, Praz CR, Sotiropoulos AG, et al (2019a) A chromosome-scale genome assembly reveals a highly dynamic effector repertoire of wheat powdery mildew. New Phytol 221:2176–2189. 10.1111/nph.15529

41. Müller MC, Praz CR, Sotiropoulos AG, et al (2019b) A chromosome-scale genome assembly reveals a highly dynamic effector repertoire of wheat powdery mildew. New Phytol 221:2176–2189. 10.1111/nph.15529

42. Neumann P, Navrátilová A, Koblížková A, et al (2011) Plant centromeric retrotransposons: A structural and cytogenetic perspective. Mob DNA 2:1–16. 10.1186/1759-8753-2-4

43. Nianiou-Obeidat I, Madesis P, Kissoudis C, et al (2017) Plant glutathione transferase-mediated stress tolerance: functions and biotechnological applications. Plant Cell Reports 2017 366 36:791–805. 10.1007/S00299-017-2139-7

44. Noh HJ, Lee YM, Park CH, et al (2020) Microbiome in Cladonia squamosa Is Vertically Stratified According to Microclimatic Conditions. Front Microbiol 11:503192. 10.3389/FMICB.2020.00268

45. Pankratov TA, Grouzdev DS, Patutina EO, et al (2020a) Lichenibacterium ramalinae gen. nov, sp. nov., Lichenibacterium minor sp. nov., the first endophytic, beta-carotene producing bacterial representatives from lichen thalli and the proposal of the new family Lichenibacteriaceae within the order Rhizobiales. Antonie van Leeuwenhoek, Int J Gen Mol Microbiol 113:477–489. 10.1007/S10482-019-01357-6

46. Pankratov TA, Grouzdev DS, Patutina EO, et al (2020b) Lichenicoccus roseus gen. Nov., sp. nov., the first bacteriochlorophyll a-containing, psychrophilic and acidophilic acetobacteraceae bacteriobiont of lichen cladonia species. Int J Syst Evol Microbiol 70:4591–4601. 10.1099/IJSEM.0.004318

47. Parks DH, Imelfort M, Skennerton CT, et al (2015) CheckM: assessing the quality of microbial genomes recovered from isolates, single cells, and metagenomes. Genome Res 25:1043–1055. 10.1101/GR.186072.114

48. Paulsen J, Allen JL, Morris N, et al (2024) Geography, Climate, and Habitat Shape the Microbiome of the Endangered Rock Gnome Lichen (Cetradonia linearis). Diversity 16:178. 10.3390/D16030178/S1

49. Pino-Bodas R, Stenroos S (2021) Global Biodiversity Patterns of the Photobionts Associated with the Genus Cladonia (Lecanorales, Ascomycota). Microb Ecol 82:173–187. 10.1007/S00248-020-01633-3

50. Puvar AC, Nathani NM, Shaikh I, et al (2020) Bacterial line of defense in Dirinaria lichen from two different ecosystems: First genomic insights of its mycobiont Dirinaria sp. GBRC AP01. Microbiol Res 233:126407. 10.1016/J.MICRES.2019.126407

51. Ramírez F, Ryan DP, Grüning B, et al (2016) deepTools2: a next generation web server for deep-sequencing data analysis. Nucleic Acids Res 44:W160–W165. 10.1093/NAR/GKW257

52. Rawat SR, Männistö MK, Starovoytov V, et al (2013) Complete genome sequence of Granulicella mallensis type strain MP5ACTX8T, an acidobacterium from tundra soil. Stand Genomic Sci 9:71–82. 10.4056/SIGS.4328031

53. Ronquist F, Teslenko M, Van Der Mark P, et al (2012) MrBayes 3.2: Efficient Bayesian Phylogenetic Inference and Model Choice Across a Large Model Space. Syst Biol 61:539–542. 10.1093/SYSBIO/SYS029

54. Roxas VP, Smith RK, Smith RK, Allen RD (1997) Overexpression of glutathione S-transferase/glutathioneperoxidase enhances the growth of transgenic tobacco seedlings during stress. Nat Biotechnol 1997 1510 15:988–991. 10.1038/nbt1097-988

55. Sanders WB (2024) The disadvantages of current proposals to redefine lichens. New Phytol 241:969–971. 10.1111/NPH.19321

56. Sharma A, Presting GG (2008) Centromeric retrotransposon lineages predate the maize/rice divergence and differ in abundance and activity. Mol Genet Genomics 279:133–147. 10.1007/S00438-007-0302-5

57. Shishido TK, Wahlsten M, Laine P, et al (2021) Article microbial communities of cladonia lichens and their biosynthetic gene clusters potentially encoding natural products. Microorganisms 9:1347. 10.3390/MICROORGANISMS9071347/S1

58. Simão FA, Waterhouse RM, Ioannidis P, et al (2015) BUSCO: assessing genome assembly and annotation completeness with single-copy orthologs. Bioinformatics 31:3210–3212. 10.1093/BIOINFORMATICS/BTV351

59. Storer J, Hubley R, Rosen J, et al (2021) The Dfam community resource of transposable element families, sequence models, and genome annotations. Mob DNA 12:. 10.1186/S13100-020-00230-Y

60. Tagirdzhanova G, Saary P, Cameron ES, et al (2024) Microbial occurrence and symbiont detection in a global sample of lichen metagenomes. PLoS Biol 22:e3002862. 10.1371/JOURNAL.PBIO.3002862

61. Trivedi P, Batista BD, Bazany KE, Singh BK (2022) Plant–microbiome interactions under a changing world: responses, consequences and perspectives. New Phytol 234:1951– 1959. 10.1111/NPH.18016

62. Tzovaras BG, Segers FHID, Bicker A, et al (2020) What Is in Umbilicaria pustulata? A Metagenomic Approach to Reconstruct the Holo-Genome of a Lichen. Genome Biol Evol 12:309–324. 10.1093/GBE/EVAA049

63. Vandenkoornhuyse P, Quaiser A, Duhamel M, et al (2015) The importance of the microbiome of the plant holobiont. New Phytol 206:1196–1206. 10.1111/NPH.13312

64. Wei T, Simko V (2021). R package ’corrplot’: Visualization of a Correlation Matrix. (Version 0.92), https://github.com/taiyun/corrplot

65. Wicaksono WA, Cernava T, Grube M, Berg G (2020) Assembly of Bacterial Genomes from the Metagenomes of Three Lichen Species. Microbiol Resour Announc 9:. 10.1128/MRA.00622-20

66. Wicker T, Stritt C, Sotiropoulos AG, et al (2022) Transposable element populations shed light on the evolutionary history of wheat and the complex co-evolution of autonomous and non-autonomous retrotransposons. Adv Genet 3:. 10.1002/GGN2.202100022

67. Wickham H, Averick M, Bryan J, et al (2019) Welcome to the Tidyverse. J Open Source Softw 4:1686. 10.21105/JOSS.01686

68. Yahr R, Vilgalys R, Depriest PT (2006) Geographic variation in algal partners of Cladonia subtenuis (Cladoniaceae) highlights the dynamic nature of a lichen symbiosis. New Phytol 171:847–860. 10.1111/J.1469-8137.2006.01792.X

