## Supplementary_Figures for "A reference metagenome sequence of the lichen *Cladonia rangiformis*"

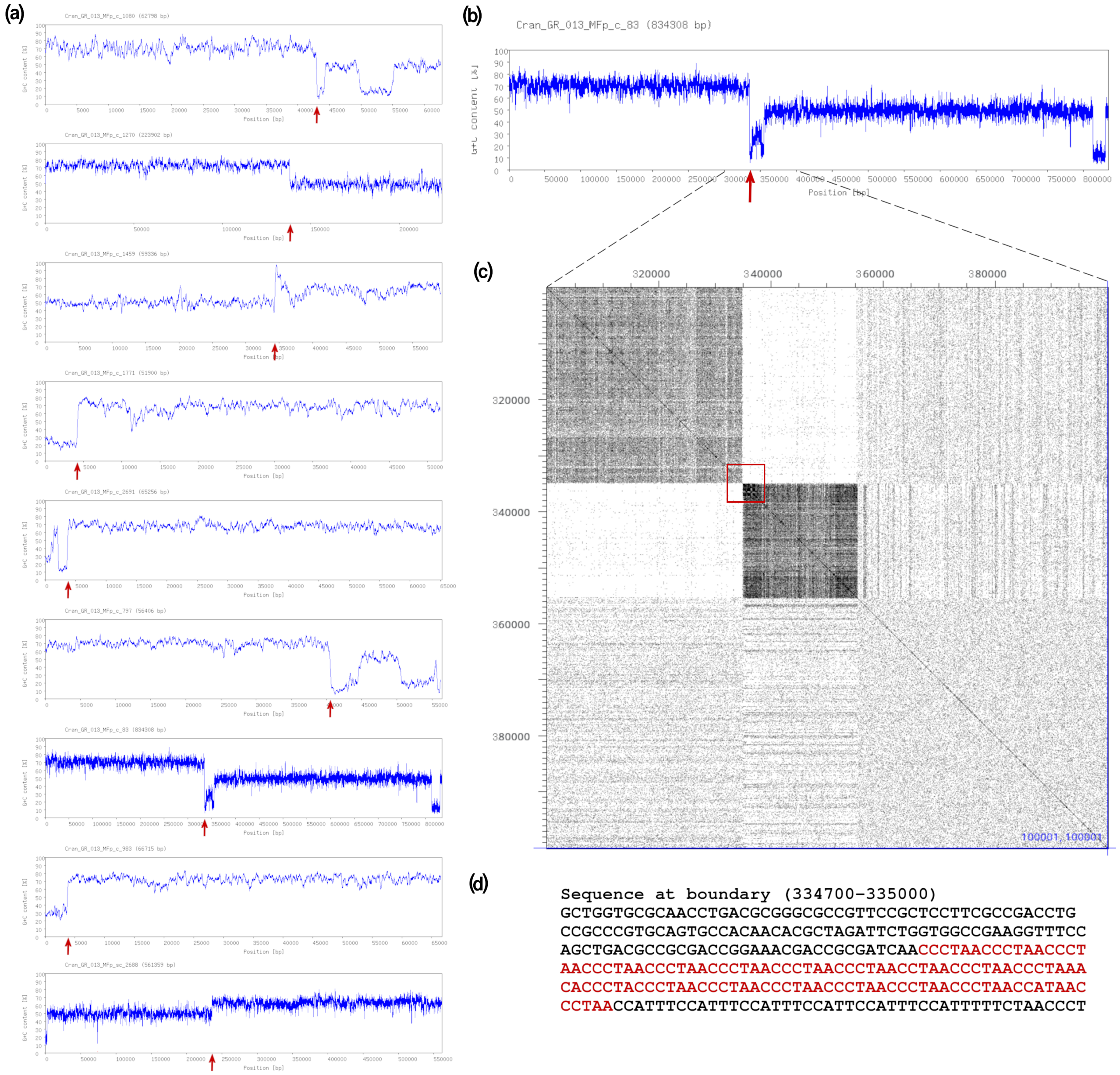


**Fig. S1** Identification and processing of chimeric sequence contigs. **(a)** Plots of CG content in 9 chimeric sequence contigs from the *C. rangiformis* reference genome assembly. GC content was calculated in sliding windows of 200 bp with a sliding step of 10 bp. The regions with lower GC content were determined to belong to the fungus while those with higher GC content were determined to belong to bacteria. The boundaries where contigs were broken up are indicated with red arrows. **(b)** GC plot of contig 83. GC content was calculated in sliding windows of 200 bp with a sliding step of 10 bp. **(c)** Dot plot alignment of the boundary region against itself. Regions of differing GC content are clearly visible due to their differing levels of background noise. The red box indicates the region to which the boundary between bacterial and fungal sequences was narrowed down. **(d)** The sequence containing the boundary. Fungal telomeric sequence repeats are shown in red.

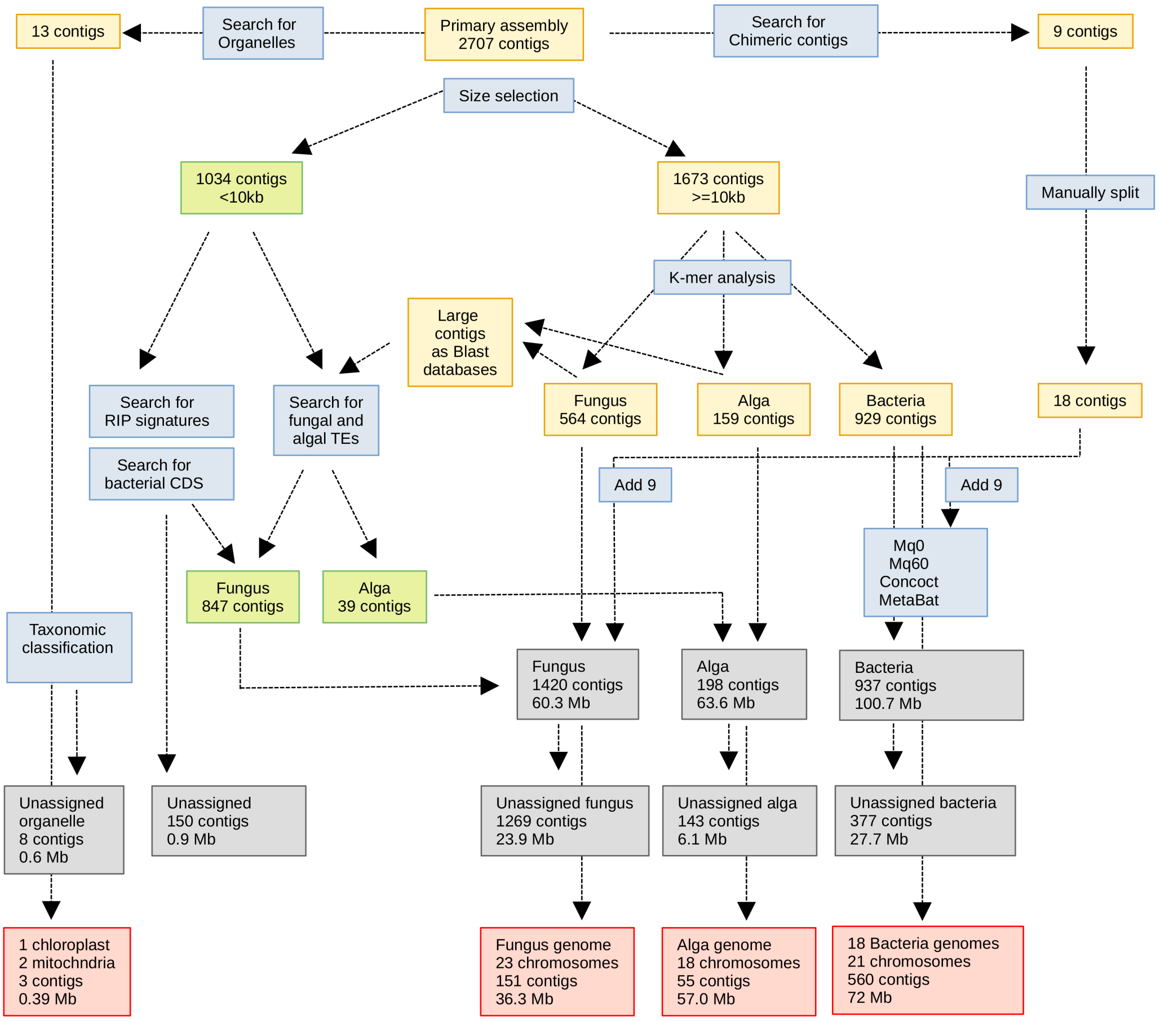


**Fig. S2** Work flow for the isolation of genomes from individual species in the reference metagenome assembly of *C. rangiformis*.

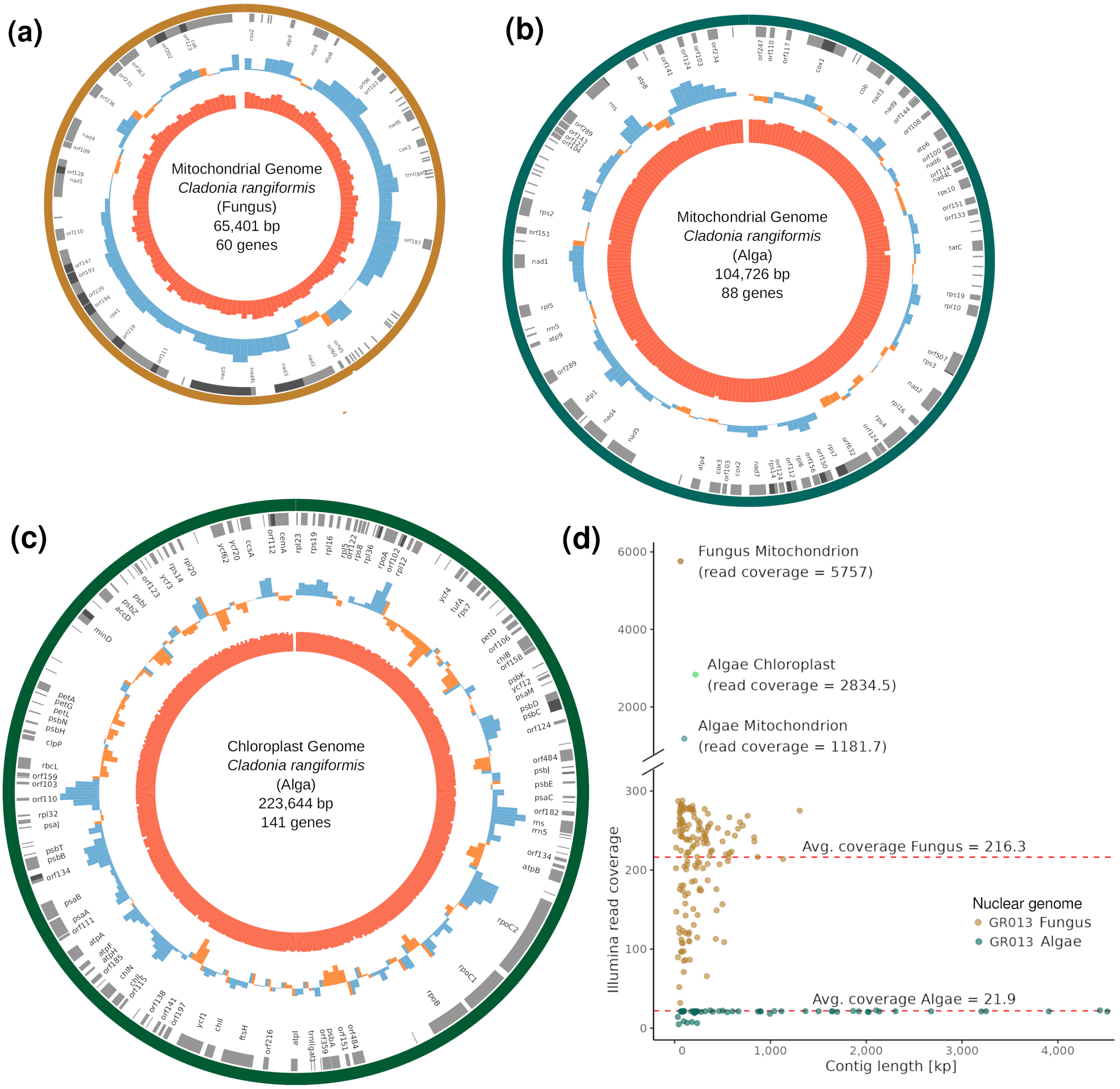


**Fig. S3** Analysis of the genomes of mitochondria from *C. rangiformis* and *A. mediterranea* and the chloroplast of *A. mediterranea.* **(a-c)** Organelle maps. The inner most circle (red) represents the GC content [%], the second circle from the center shows the GC skew (in blue positive, in orange negative), the outer circle (varying colours depending on the organelle-species) depicts the genes. The genes in darker gray are either overlapping with or directly adjacent to one another. **(a)** *C. rangiformis* mitochondrium genome. **(b)** *A. mediterranea* mitochondrium genome. **(c)** *A. mediterranea* chloroplast genome. **(d)** Ilumina read coverage of PacBio sequence contigs from the organellar and nuclear genomes. Shown are 151 and 56 individual contigs that were used for the chromosome-scale assembly of the fungal and algal nuclear genomes, respectively, as well as the three contigs that represent the organelles. Average sequence coverage of fungal and algal nuclear genomes is indicated with dashed red lines. From these values, we estimate that there are approximately 26.6 (5757/216.3) mitochondrial genomes per fungal nuclear genome. Analogously, we estimate that per algal nuclear genome, there are ~56 (1181.7/21.9) mitochondrial and ~134 (2834.5/21.9) chloroplast genomes.

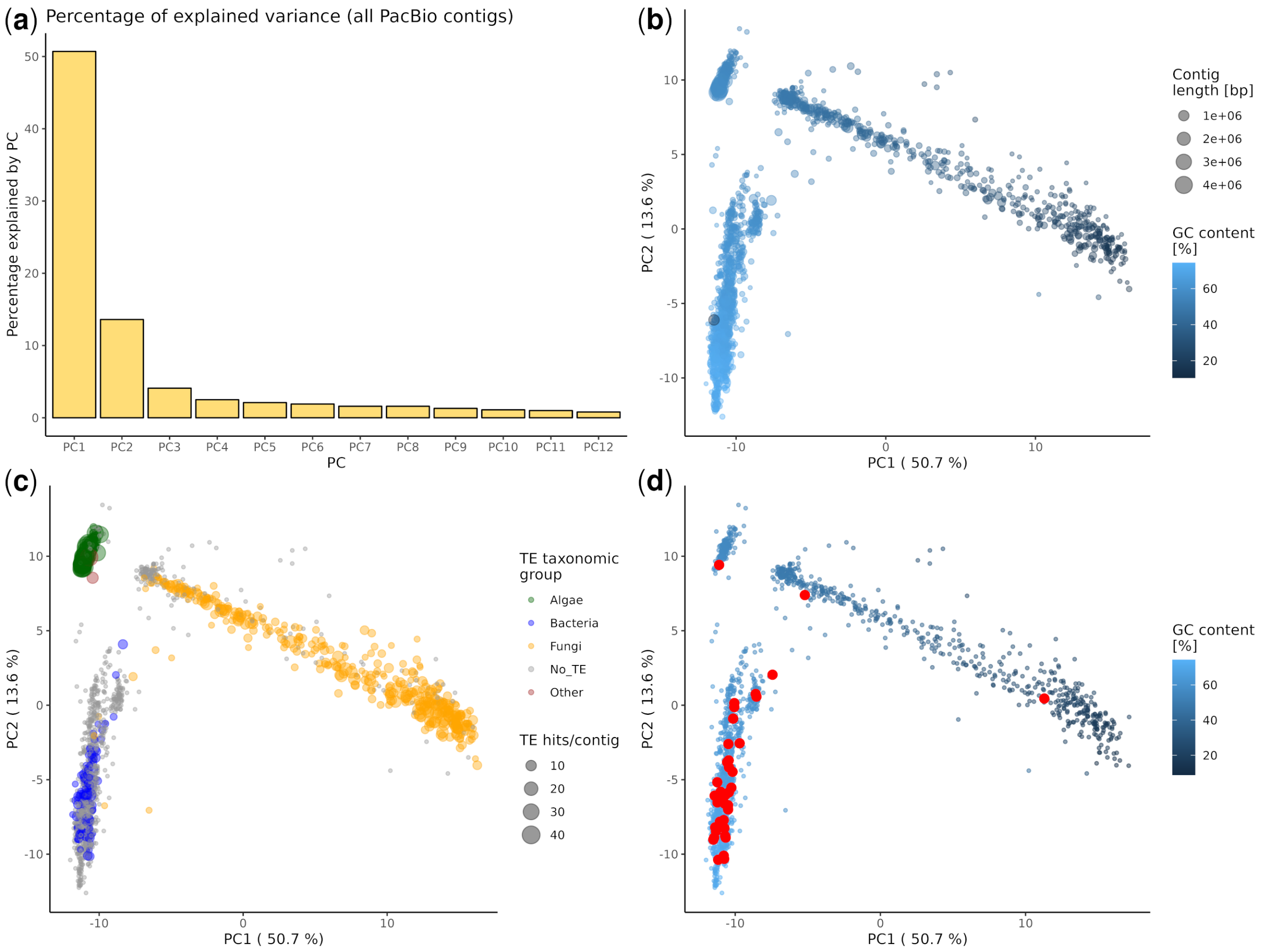


**Fig. S4** Principal component analysis (PCA) of k-mer frequencies in 1675 sequence contigs longer than 10 kb form the *C. rangiformis* reference metagenome assembly. Frequencies of all 256 possible tetra-nucleotides were calculated for all individual contigs. **(a)** Histogram of the percentages explained by PC1 through PC12. **(b)** PCA of 1675 sequence contigs colored by GC content. **(c)** PCA of 1675 sequence contigs with superimposed blastx hits to predicted TE proteins. Hits to TE proteins allows confident classification into three main organism groups, alga, fungus and bacteria. **(d)** PCA of 1675 sequence contigs with superimposed in red the contigs on which ribosomal gene clusters were identified.

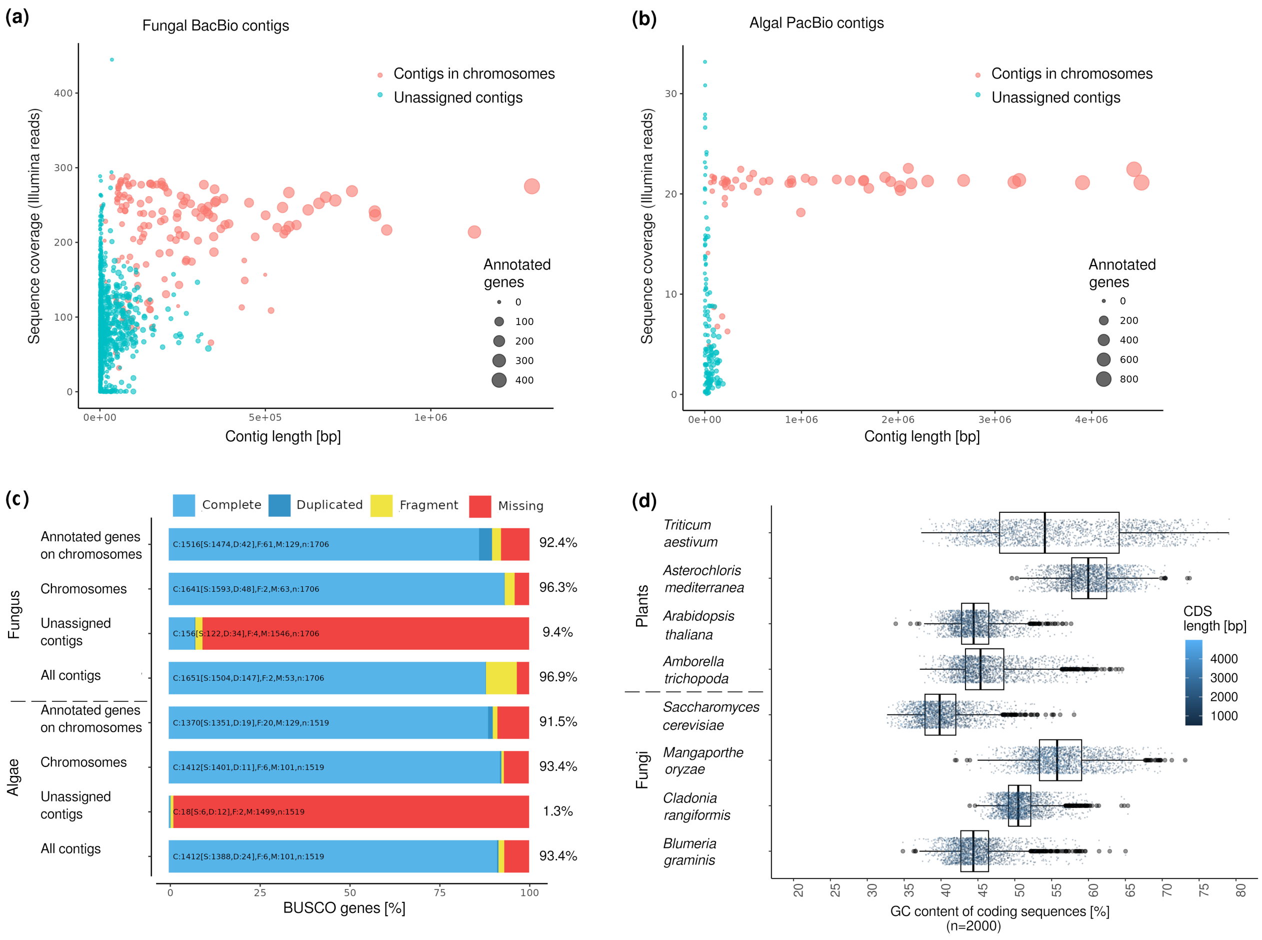


**Fig. S5** Chromosome assembly and Analysis of gene content of *C. rangiformis* (fungus) and *A. mediterranea* (algae) genomes. **(a)** Length and read coverage of PacBio contigs that were classified as coming from fungi. In red are contigs that were used to assemble the chromosome-scale scaffolds with bubble size indicating the number of annotated genes. **(b)** The same plot for PacBio contigs that were classified as coming from algae. **(c)** BUSCO scores of fungal and algal genome using the datasets ascomycota_odb10 and chlorophyta_odb10, respectively. For both genomes scores are shown for contigs that were assembled into chromosomes, unassigned contigs and all contigs. Additionally, the top bar shows BUSCO on genes annotated in chromosomes. The numbers to the right of the bars indicated the total for complete, duplicated and fragmented genes. Note that unassigned contigs have very low BUSCO values, indicating that they mostly represent repetitive sequences that were assembled from species/haplotypes that were present in much lower abundance than the main fungus and alga. **(d)** Comparison of GC content of predicted coding sequences (CDS) from different species. For each species 2,000 randomly picked genes were used. Each dot represents one gene with the color indicating the length of the coding sequence (CDS). Boxes indicate the inter-quartile range (IQR) with the central line indicating the median and whiskers indicating the minimum and maximum without outliers, respectively. Outliers were defined as minimum – 1.5*IQR and maximum + 1.5*IQR, respectively.

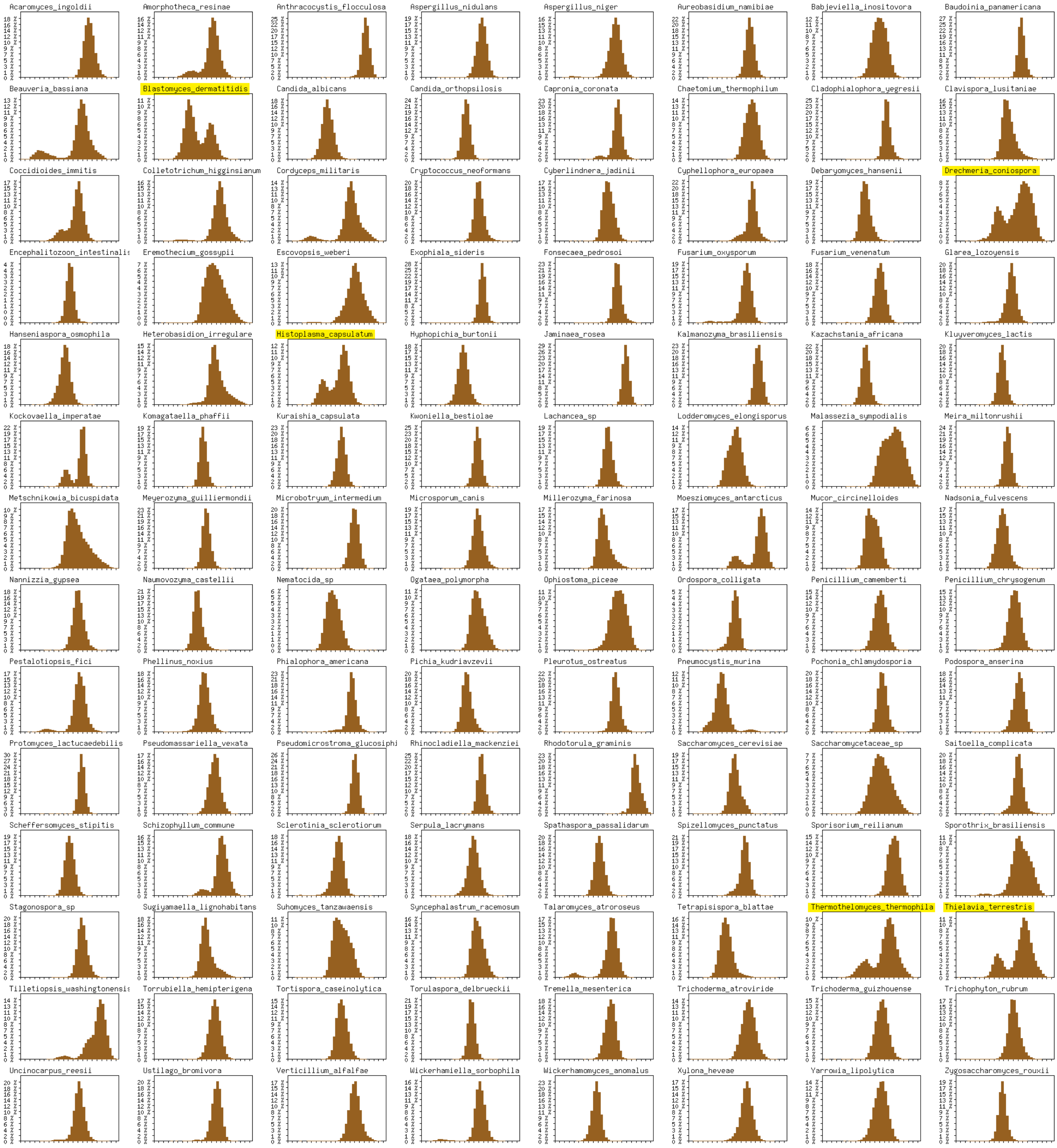

**Fig. S6** GC content profiles from 112 ascomycete genomes. The genomes were split into non-overlapping 500 bp segments for which GC content was calculated. The x axis indicates the GC content [%], while the y axis indicates the number of segments with a given GC content. Highlighted are those genomes which show a a bimodal GC content distribution similar to that of *C. rangiformis*.

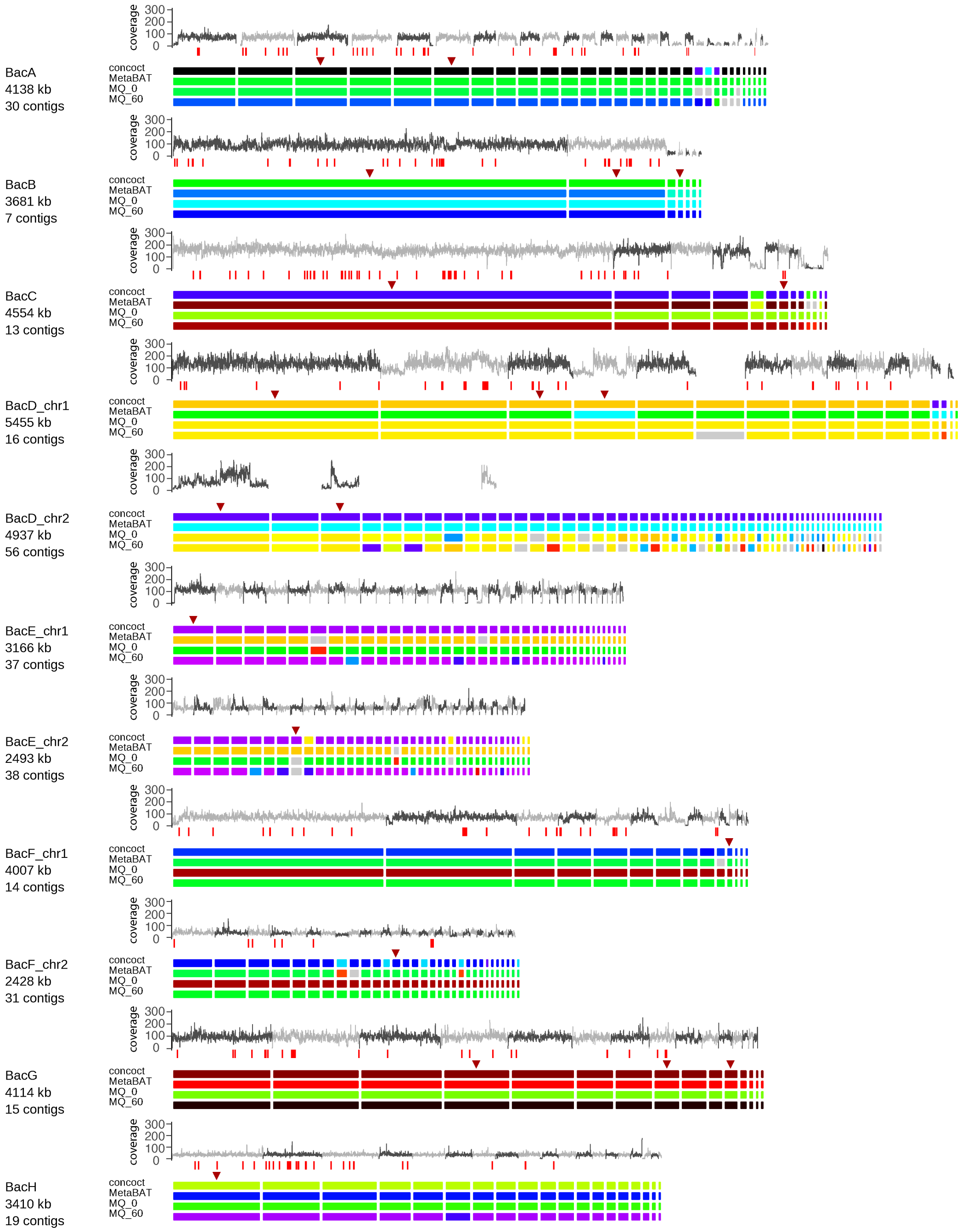


**Supplementary Fig. S7 part 1.** Binning of PacBio sequence contigs into putative bacterial genomes. The top track shows the sequence coverage with illumina reads in windows of 1000 bp. Underneath are positions of BUSCO genes indicated as red vertical bars. The 4 coloured tracks indicate the bins in which sequence contigs were placed by the four binning methods. Triangles indicate the sequence contigs on which 16S ribosomal DNA (rDNA) loci were found. These were used for taxonomic classification of bacteria if the respective contig was binned into the largest group by at least 3 binning methods. Note that some of the contigs do not show coverage, because of the way bamCoverage merges bins with low coverage.

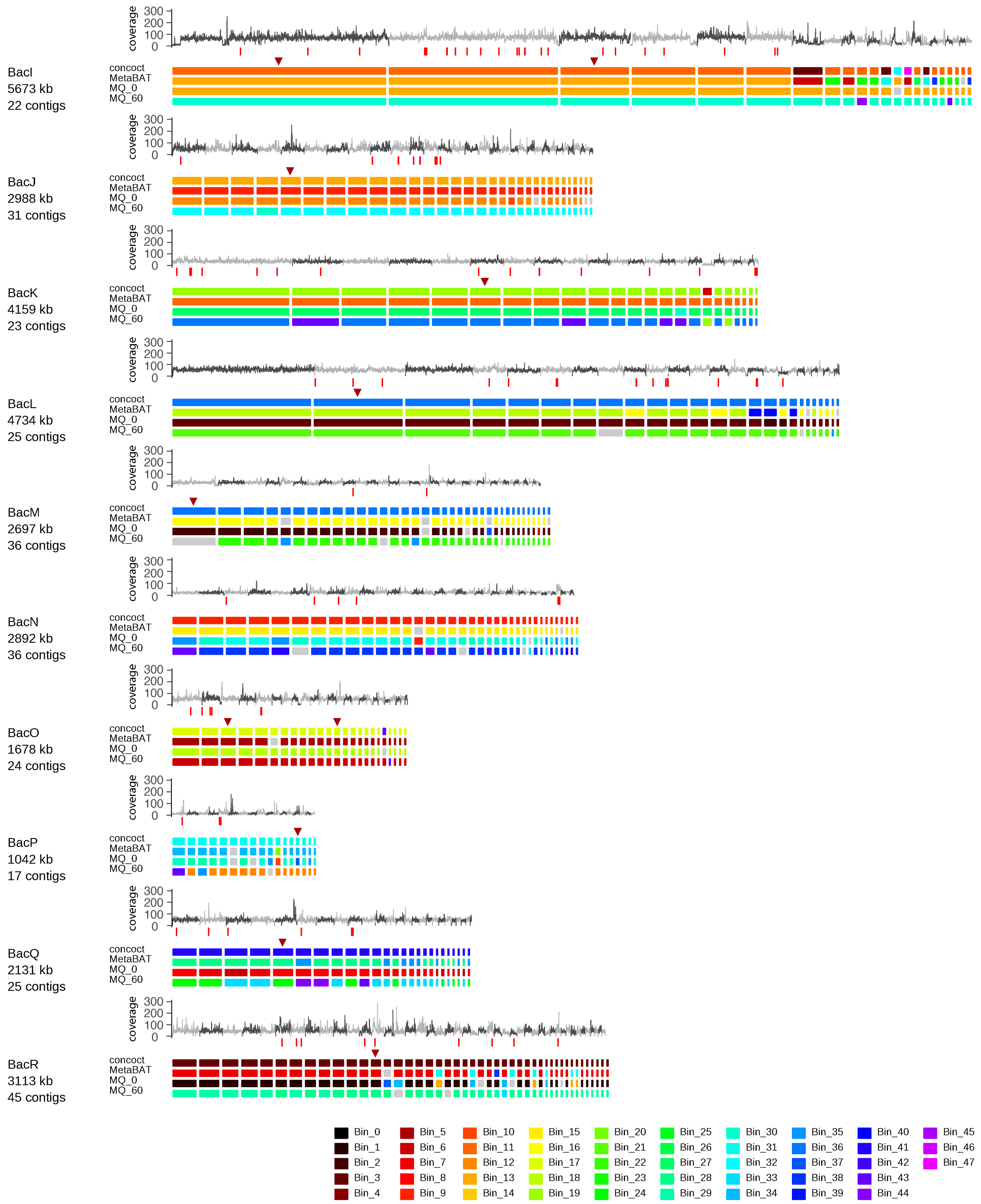


**Supplementary Fig. S7 (continued).** Binning of PacBio sequence contigs into putative bacterial genomes. The top track shows the sequence coverage with illumina reads in windows of 1000 bp. Underneath are positions of BUSCO genes indicated as red vertical bars. The 4 coloured tracks indicate the bins in which sequence contigs were placed by the four binning methods. Triangles indicate the sequence contigs on which 16S ribosomal DNA (rDNA) loci were found. These were used for taxonomic classification of bacteria if the respective contig was binned into the largest group by at least 3 binning methods.

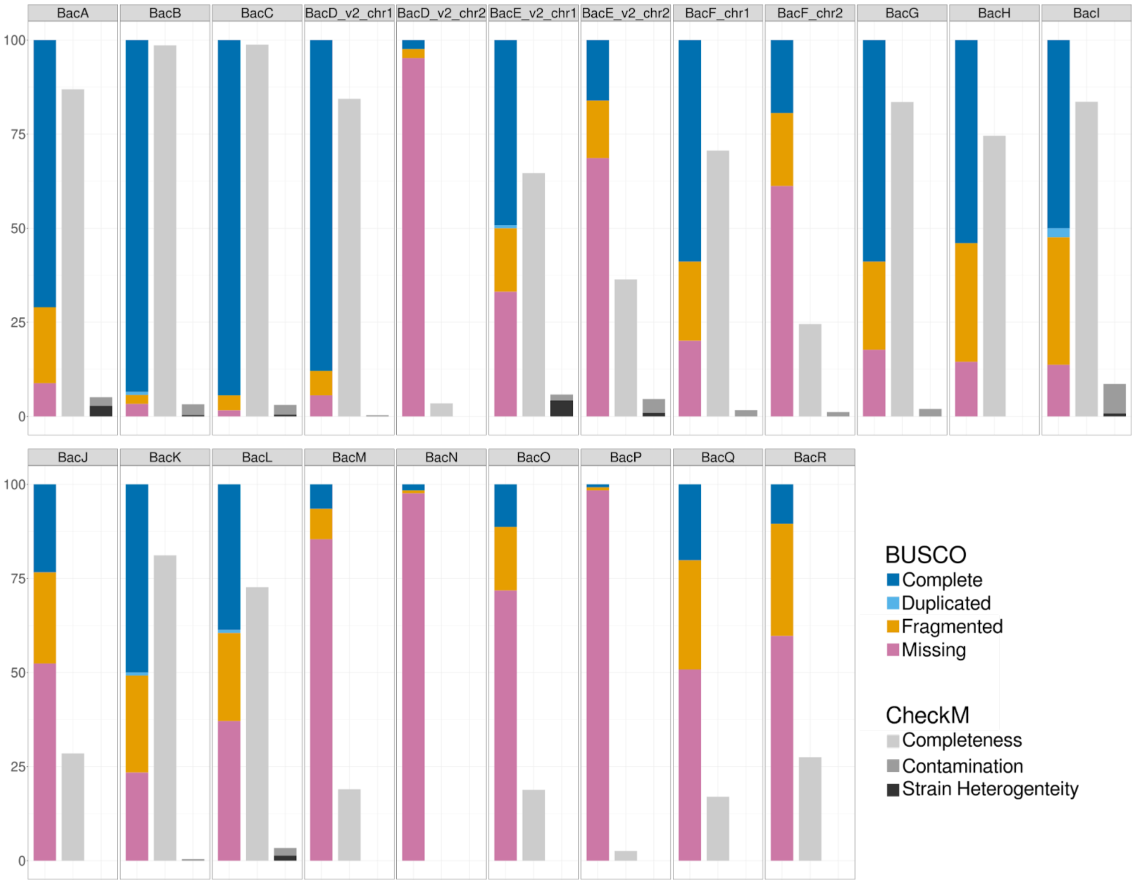


**Fig. S8** BUSCO and CheckM scores of bacterial genomes. The first bar represents the BUSCO scores, the second and third bar display the CheckM scores for completeness and contamination with strain heterogeneity respectively. For the Complete values (dark blue) shown in the figure the raw duplicated BUSCO score (light blue) was subtracted from the raw complete values of BUSCO. Thus, the total BUSCO complete value is composed of the Complete and Duplicated section displayed in the figure. The same approach was taken for the CheckM Contamination and Strain Heterogeneity. The total Contamination value from CheckM is composed of the Contamination and Strain Heterogeneity shown in the figure.

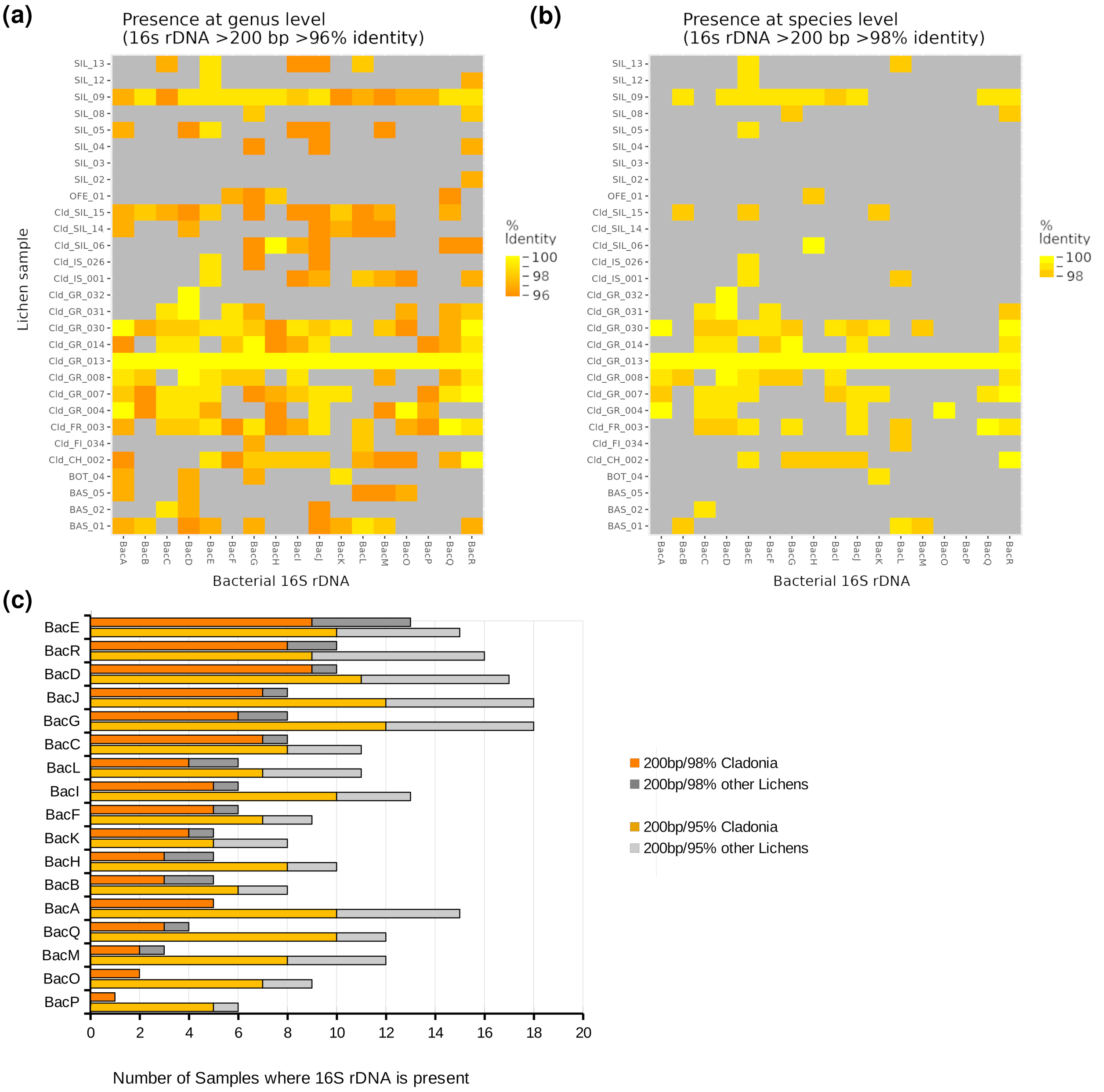


**Fig S9.** Test for presence of the bacteria identified in C. rangiformis in 29 lichen samples. Samples with the prefix "Cld_" come from lichens of the genus *Cladonia*. **(a)** Graphical summary of blastn hits of 16S rDNA in short read assemblies of 29 lichen samples. Here, it was examined whether a 16S rRNA of an bacteria species identified in the *C. rangiformis* (Cld_GR_013) sample has blast hits with >95% sequence identify in any of the other samples. The short minimal hit length of 200 was chosen to accommodate for the fact that many Illumina short read assemblies are highly fragmented. If a sample contained a sequence with >95% identity, it was assumed that it contained bacteria of the same genus as the one from the *C. rangiformis* (Cld_GR_013) sample. **(b)** analogous analysis as in **(a)** with a threshold of 98%, identifying bacteria of the same species. **(c)** Numbers of blastn hits for 16S rDNA of the identified bacteria in the 29 lichen samples. Counts were made for lichens of the Cladonia genus (orange) and other lichens (gray). Count are given for sequence identity >=98% (for same species) and >=95% (for same genus).

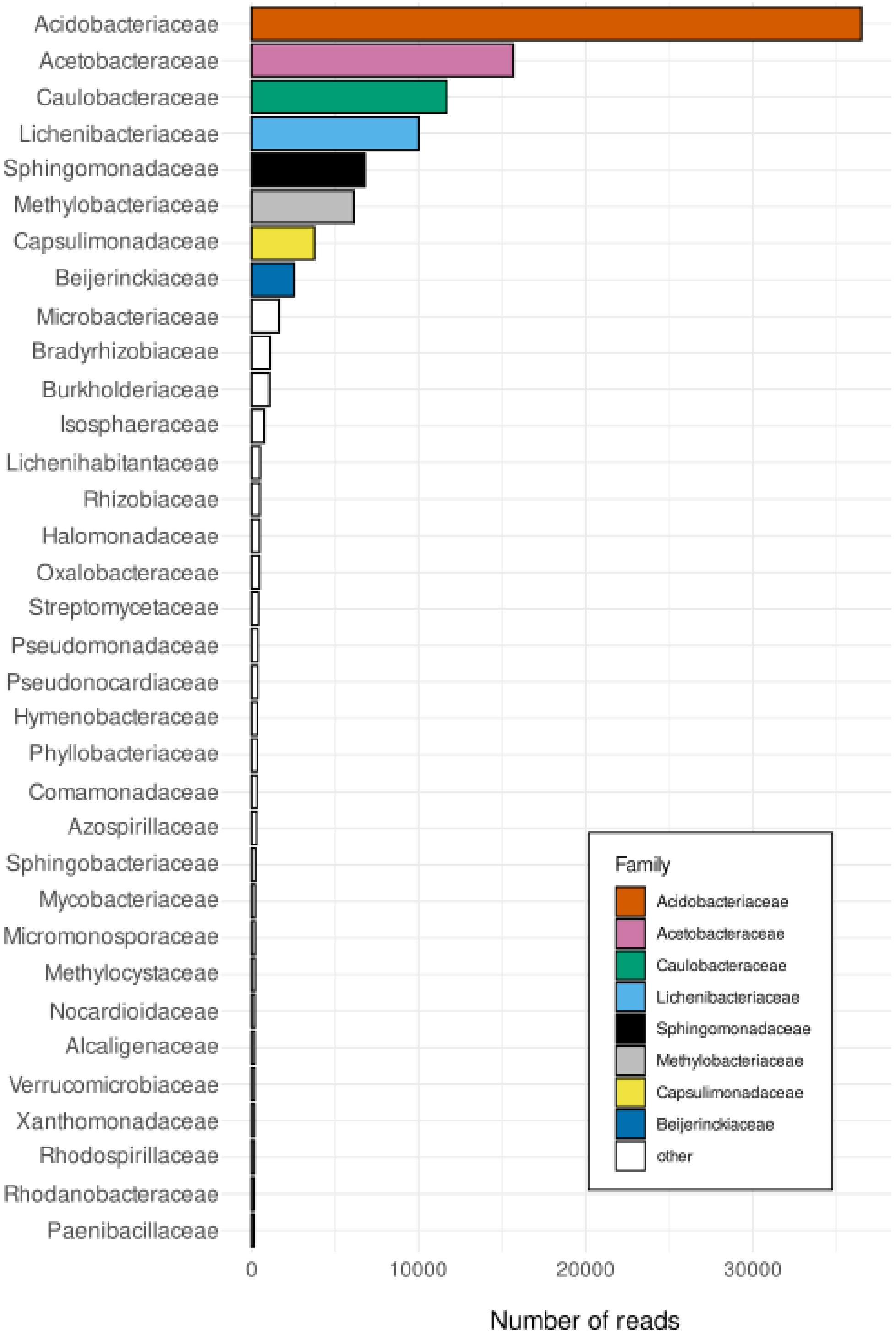


**Fig. S10** MEGAN classified illumina reads wich could be classified to the family level in the GR013 lichen sample. The top eight families taken together make up ~90% (93096 of 104093 reads) of the reads that could be classified at least to bacteria family level.

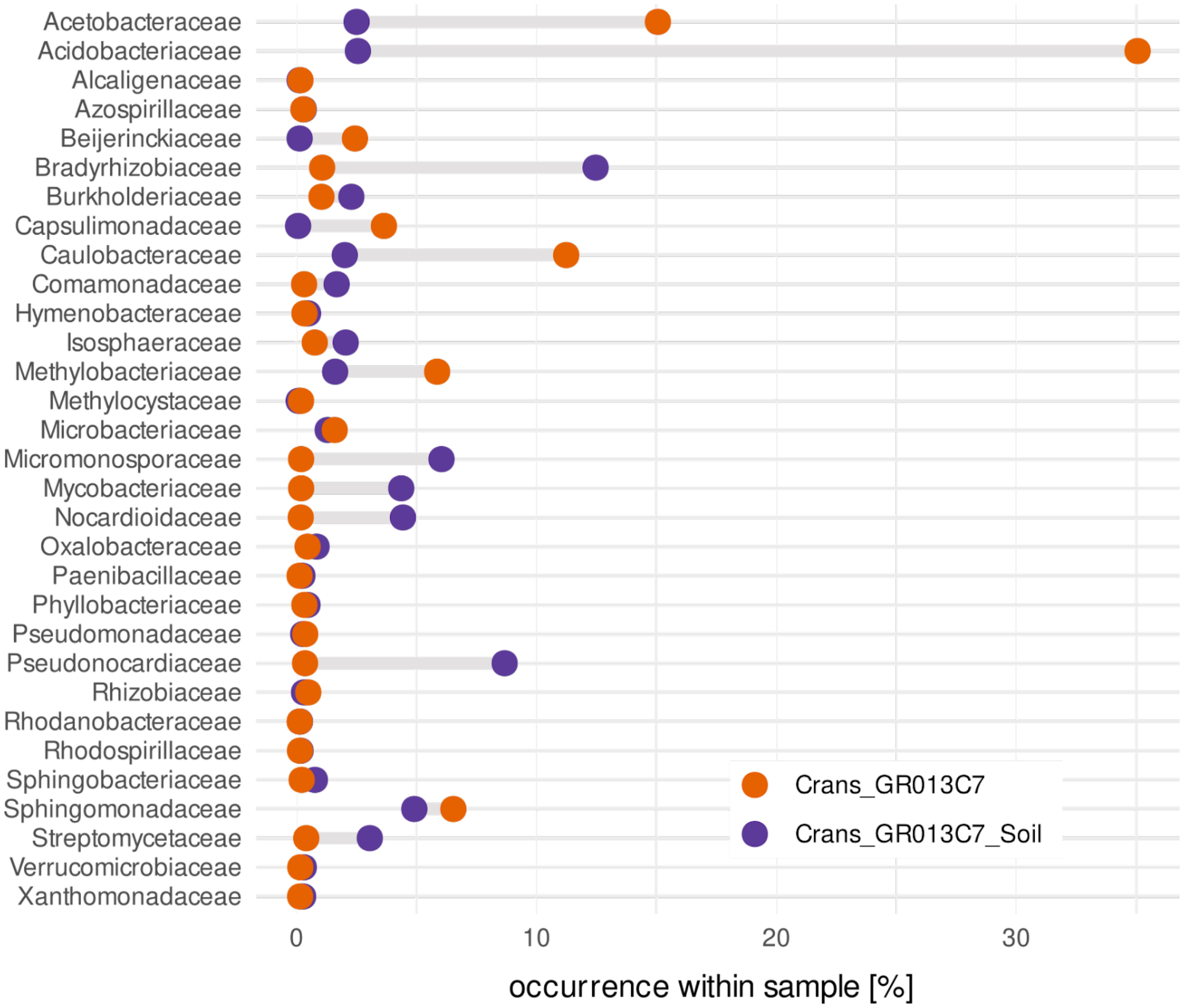


**Fig. S11** Proportional occurrence of bacterial families common to *C. rangiformis* lichen and corresponding soil samples.

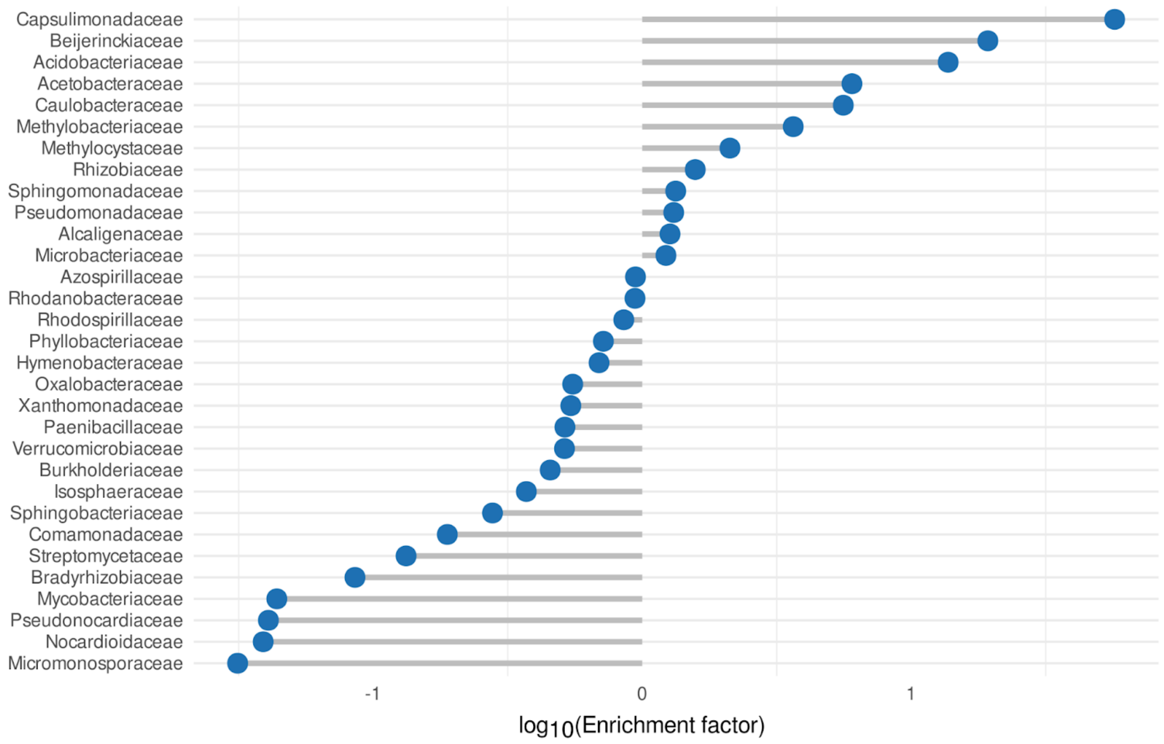


**Fig. S12** Enrichment factor of bacterial families found in both, the *C. rangiformis* lichen and its corresponding soil samples. Negative values indicate enrichment in the soil sample while positive values indicate enrichment in the lichen sample.

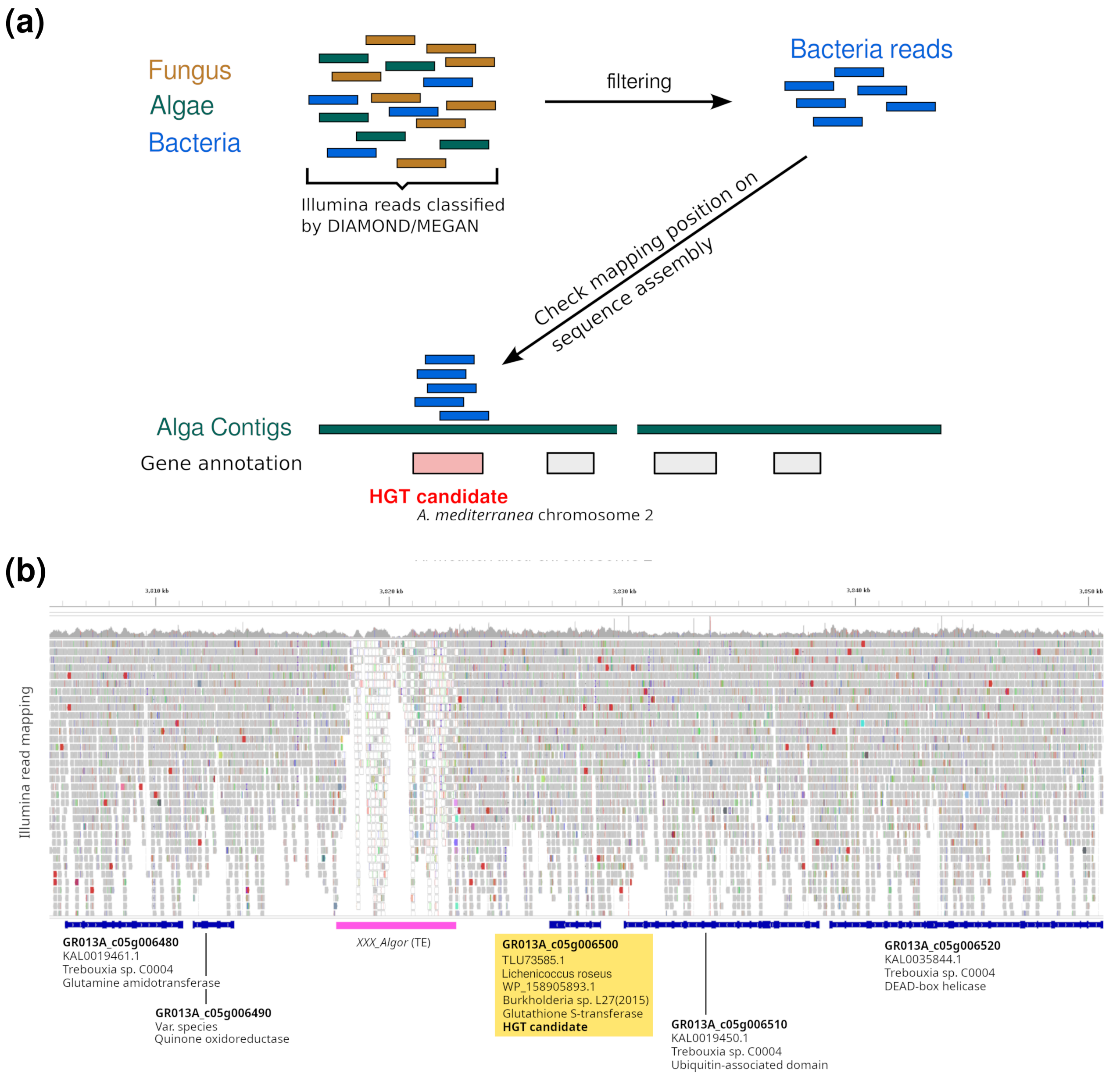


**Fig. S13.** Identification of putative horizontal gene transfers (HGTs). **(a)** Schematic workflow for initial HT candidate identification. We identified the candidate proteins by using read classification from MEGAN and filtered mapping information. Here, we compiled raw read classification with information from read mapping to the metagenome assembly. In the schematic shown, we first filtered fro reads that were classified by the DIAMOND/MEGAN pipeline as coming from Bacteria. Then, we analyzed where these reads map on sequence contigs from the alga *A. mediterranea*. and searched for loci where multiple reads map. It such a region harbored an annotated gene, we considered it a candidate for HGT. **(b)** Visualization of Ilumina read mapping by IGV software (igv.org) in the ~45 kb region on chromosome 2 harboring the HGT candidate gene *GR013A_c05g006500*. The top track shows coverage with genomic illumina reads from *C. mediterranea* mapped on the reference genome and individual read mappings, respectively. The bottom track shows gene annotation with exons as wider and introns as narrow bars. Note that the region has no sequence gaps and shows a largely even read coverage, excluding the possibility of a mis-assembly. The region with ambiguous read mappings to the left of the HGT candidate is due to the presence of a multi-copy TE (*XXX_Algor*). Description underneath genes indicate similarity to closest homologs in blastp searches against NCBI. Three of the neighbors of the HGT candidate all have closest hits to proteins from unicellular algae.

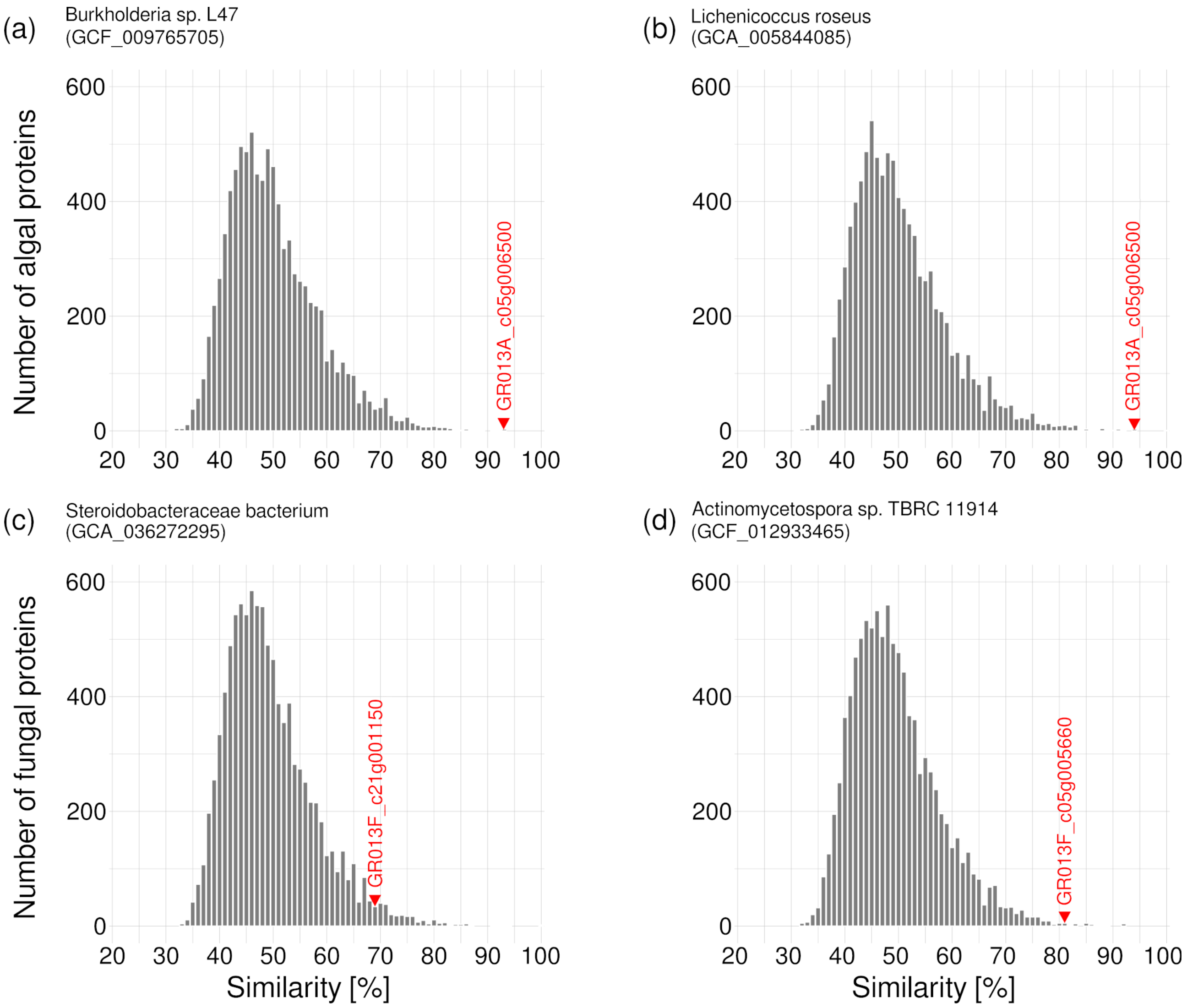


**Fig. S14** Distributions of sequence similarity [%] for algal (a and b) and fungal proteins (c and d) against bacterial proteins. The bacteria species were chosen based on the top blast hit of HGT candidates on a bacteria species. The position of the HGT candidates in the distribution is indicated with a red arrow head and the name of the gene idnetifier. **(a)** Sequence similarity of *A. mediterranea* proteins against proteins in the *Burkholderia* species that produced the top blast hit. The data for the plot was generated by using all *A. mediterranea* proteins as queries in blastp searches against the proteins encoded by the *Burkholderia* genome (NCBI accession GCF_009765705). Note that the HGT candidate GR013A_c05g006500 is among the proteins with highest similarity to the Brukholderia proteins. **(b)** The same analysis with *A. mediterranea* proteins used in blast searches against *Lichenicoccus roseus* proteins, the bacteria species that produced a blast hit nearly as strong as Burholderia shown in **(a)**. **(c)** The same analysis for *C. rangiformis* proteins used in blast searches against proteins form a bacterium of the *Steroidobacteriaceae* family. **(d)** The same analysis for *C. rangiformis* proteins used in blast searches against proteins form a protein from the *Actinomycetospora*

genus. Note that the HGT candidates are at the very extremes of the distribution, meaning that they are among those which encode proteins with the highest similarity to bacterial proteins.

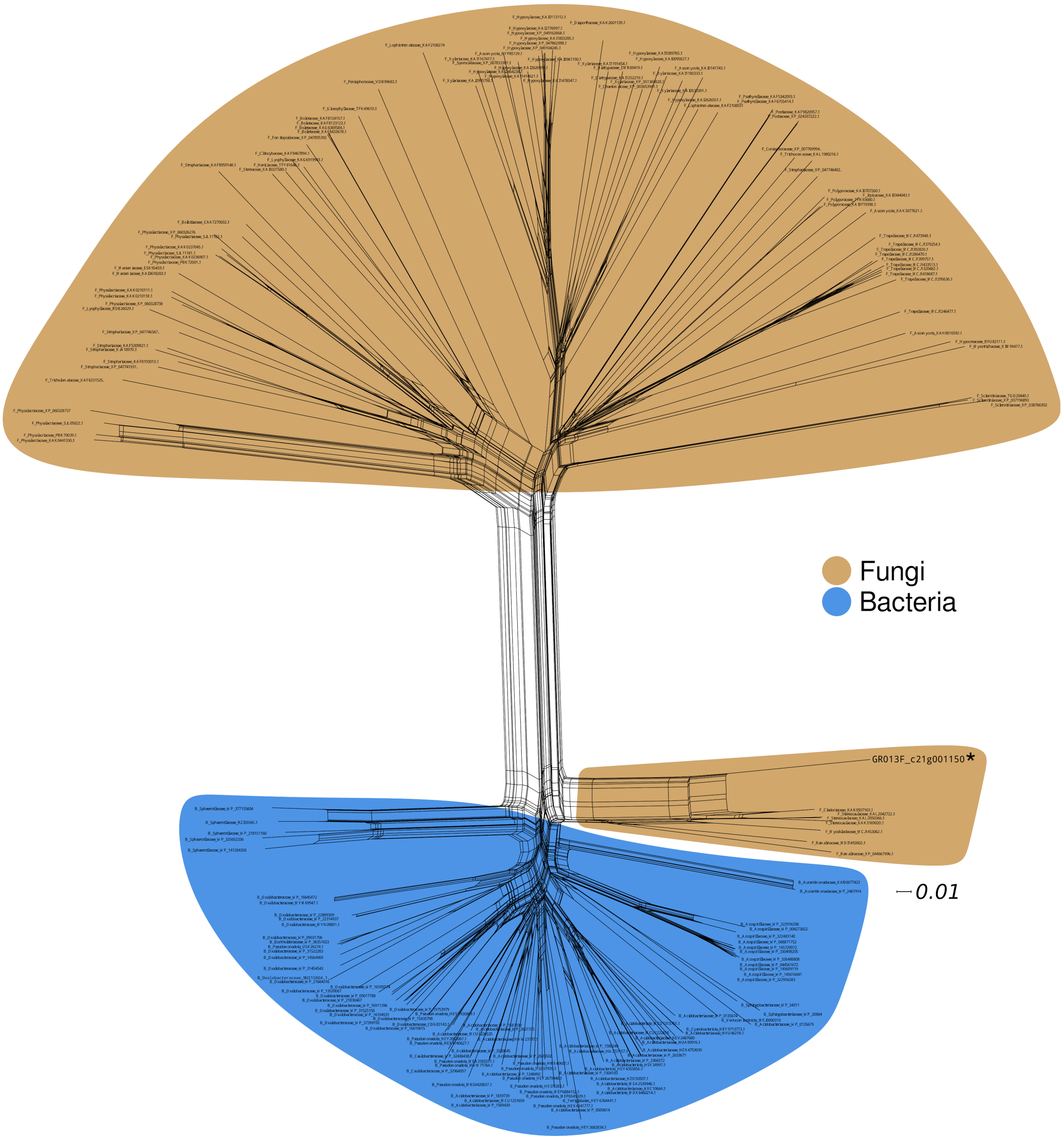


**Figure S15.** Network of proteins from the NCBI database with the horizontal gene transfer candidate *GR013F_c21g001150*. The top 100 blastp hits against the entire NCBI protein database and the top 100 blastp hits against the Fungi database (taxid:4751) were and combined with the candidate protein to generate a NeighborNet network utilizing Splitstree4. The HGT candidate is indicated in larger font and marked with an asterisk. The scale bar indicates the distance derived from the multiple alignment. Some family names had to be shortened for the labels.

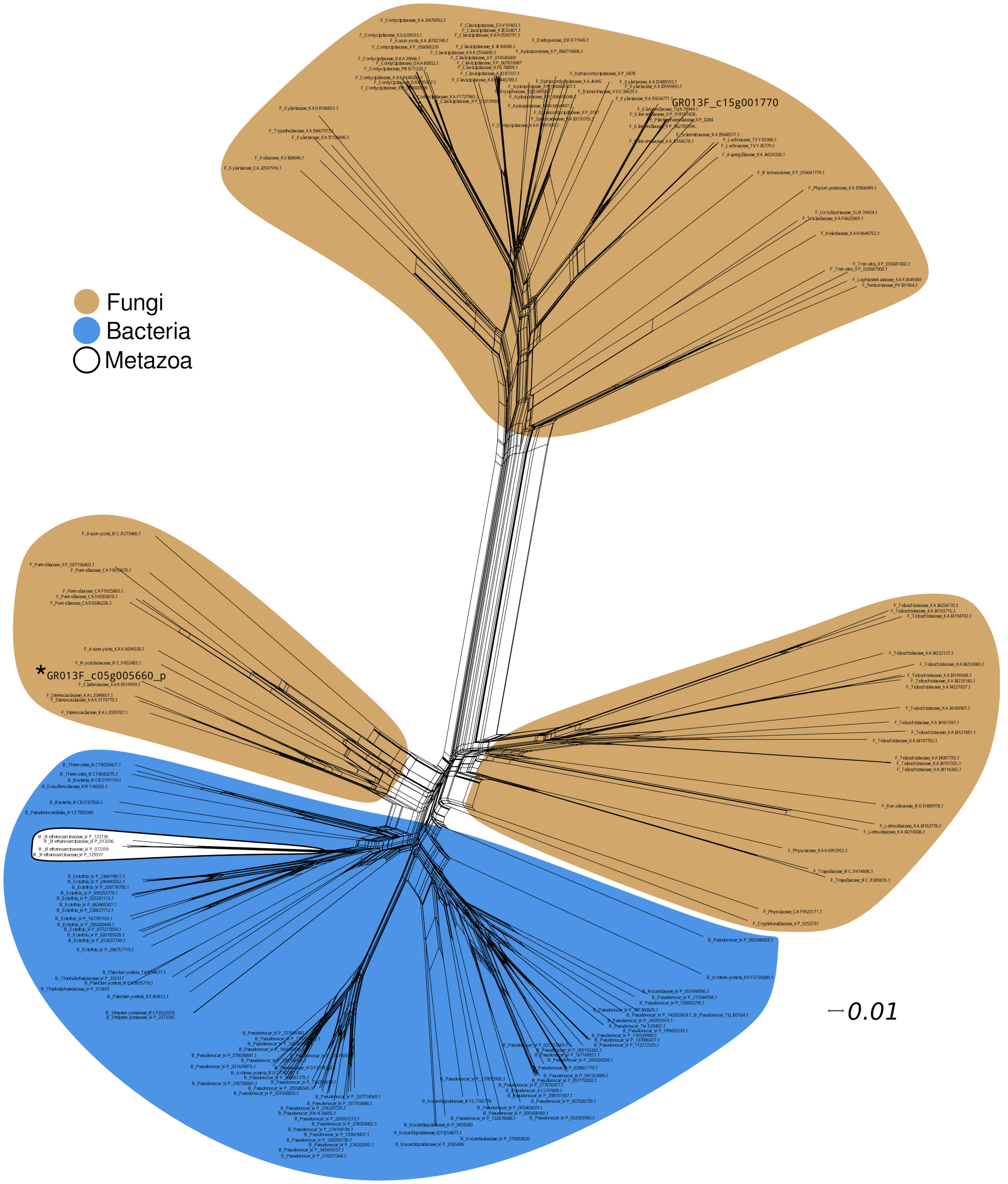


**Figure S16.** Network of proteins from the NCBI database with the horizontal gene transfer candidate *GR013F_c05g005660*. The top 100 blastp hits against the entire NCBI protein database and the top 100 blastp hits against the Fungi database (taxid:4751) were and combined with the candidate protein to generate a NeighborNet network utilizing Splitstree4. The HGT candidate is indicated in larger font and marked with an asterisk. The scale bar indicates the distance derived from the multiple alignment. Some family names had to be shortened for the labels.
